# Effect of allelopathy on plant performance

**DOI:** 10.1101/2020.05.14.095190

**Authors:** Zhijie Zhang, Yanjie Liu, Ling Yuan, Ewald Weber, Mark van Kleunen

## Abstract

Allelopathy (i.e. chemical interactions between plants) is known to affect individual performance, community structure and plant invasions. Yet, a quantitative synthesis is lacking. We performed a meta-analysis of 385 studies that measured allelopathic effects of one species (allelopathy plant) on another species or itself (test plant). Overall, allelopathy reduced plant performance by 25%, but the variation in allelopathy was high. Type of method affected allelopathic effect. Compared to leachates, allelopathy was more negative when residues of allelopathy plants were applied, and less negative when soil conditioned by allelopathy plants was applied. The negative effects of allelopathy diminished with study duration, and increased with concentrations of leachates or residues. Although allelopathy was not significantly related to life span, life form and domestication of the interacting plants, it became more negative with increasing phylogenetic distance. Moreover, native plants suffered more negative effects from leachates of naturalized alien plants than of other native plants. Our synthesis reveals that allelopathy could contribute to success of alien plants. The negative relationship between phylogenetic distance and allelopathy indicates that allelopathy might drive coexistence of close-related species (i.e. convergence) or dominance of single species.

## Introduction

Allelopathy refers to either the inhibitory or stimulatory effect of one plant on another through the production of chemicals and their release into the environment^1,2^. The earliest written record of allelopathy can be traced back to *c.* 300 BC, when the Greek philosopher Theophrastus noticed that, chick pea plants ‘exhaust’ the soil and destroy weeds^3^. To understand the roles of allelopathy in agriculture, forestry and old-field succession, numerous studies have been conducted since 1970^2,4^. In the late-1990s, research on allelopathy gained even more attention as some studies revealed allelopathy as a driving mechanism for the success of some invasive plants^5–7^. Despite the long-standing interest and the mounting number of studies on allelopathy, we still lack a quantitative synthesis of those studies. A quantitative synthesis, such as a meta-analysis^8^, is important because it allows estimation of the overall effect of allelopathy, and to assess what may cause variation in allelopathy. In other words, we could assess whether allelopathy generally inhibits or stimulates plants, and under which conditions. Additionally, the past decade has witnessed a slowdown of research on allelopathy (Fig. S1), which might result from the fact that most hypotheses on allelopathy have been tested already many times or that some of the techniques to study allelopathic effects have been questioned (e.g. Blair et al.^9^). A quantitative synthesis is needed to identify knowledge gaps and to guide future research.

The results of studies on allelopathy, like for any other topic in ecology and evolution, are heterogeneous^10^. Negative effects of allelopathy have been found in some studies^11^, but no or positive effects in others^12,13^. Identifying the sources of this heterogeneity can improve our understanding of allelopathy^14–16^. The first candidate source of heterogeneity is the design of studies. For example, to test allelopathy, some studies soaked seeds in leaf leachates or root exudate from other plants, and some grew seedlings on substrate mixed with residues from other plants (see Table 1 for a summary of the major methods that have been used to test allelopathy). The differences in study design partly reflect the four pathways through which allelochemicals are released into the environment, that is, leaching from plants by rain, decomposition of plant residues (e.g. litter), exudation from roots and volatilization (Fig. 2; Rice^2^ p309). However, studies also differ with regard to the different life stages of plants (e.g. germination *vs.* growth) and the duration of the experiment (Fig. S2). Disentangling the importance of the different factors related to the study design can hardly be achieved in a single case study, but is feasible in a meta-analysis.

**Table 1.**
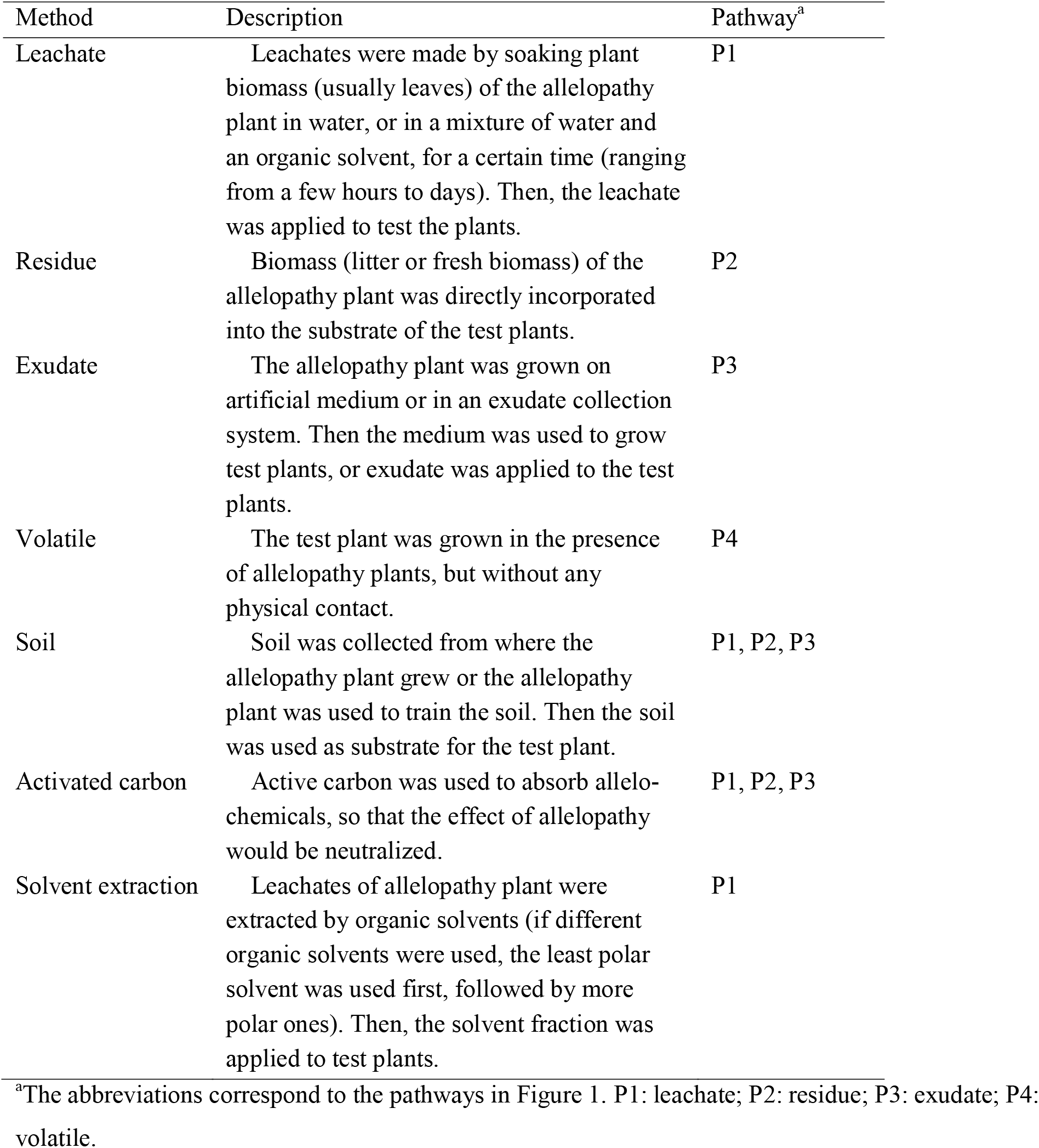
A description of the seven types of methods used to test for allelopathy. We also indicate which of the four pathways of allelopathy (see Fig. 1) the method tests.

| Method | Description | Pathway<sup>a</sup> |
| --- | --- | --- |
| Leachate | Leachates were made by soaking plant biomass (usually leaves) of the allelopathy plant in water, or in a mixture of water and an organic solvent, for a certain time (ranging from a few hours to days). Then, the leachate was applied to test the plants. | P1 |
| Residue | Biomass (litter or fresh biomass) of the allelopathy plant was directly incorporated into the substrate of the test plants. | P2 |
| Exudate | The allelopathy plant was grown on artificial medium or in an exudate collection system. Then the medium was used to grow test plants, or exudate was applied to the test plants. | P3 |
| Volatile | The test plant was grown in the presence of allelopathy plants, but without any physical contact. | P4 |
| Soil | Soil was collected from where the allelopathy plant grew or the allelopathy plant was used to train the soil. Then the soil was used as substrate for the test plant. | P1, P2, P3 |
| Activated carbon | Active carbon was used to absorb allelochemicals, so that the effect of allelopathy would be neutralized. | P1, P2, P3 |
| Solvent extraction | Leachates of allelopathy plant were extracted by organic solvents (if different organic solvents were used, the least polar solvent was used first, followed by more polar ones). Then, the solvent fraction was applied to test plants. | P1 |

<sup>a</sup>The abbreviations correspond to the pathways in Figure 1. P1: leachate; P2: residue; P3: exudate; P4: volatile.

**Figure 1.**
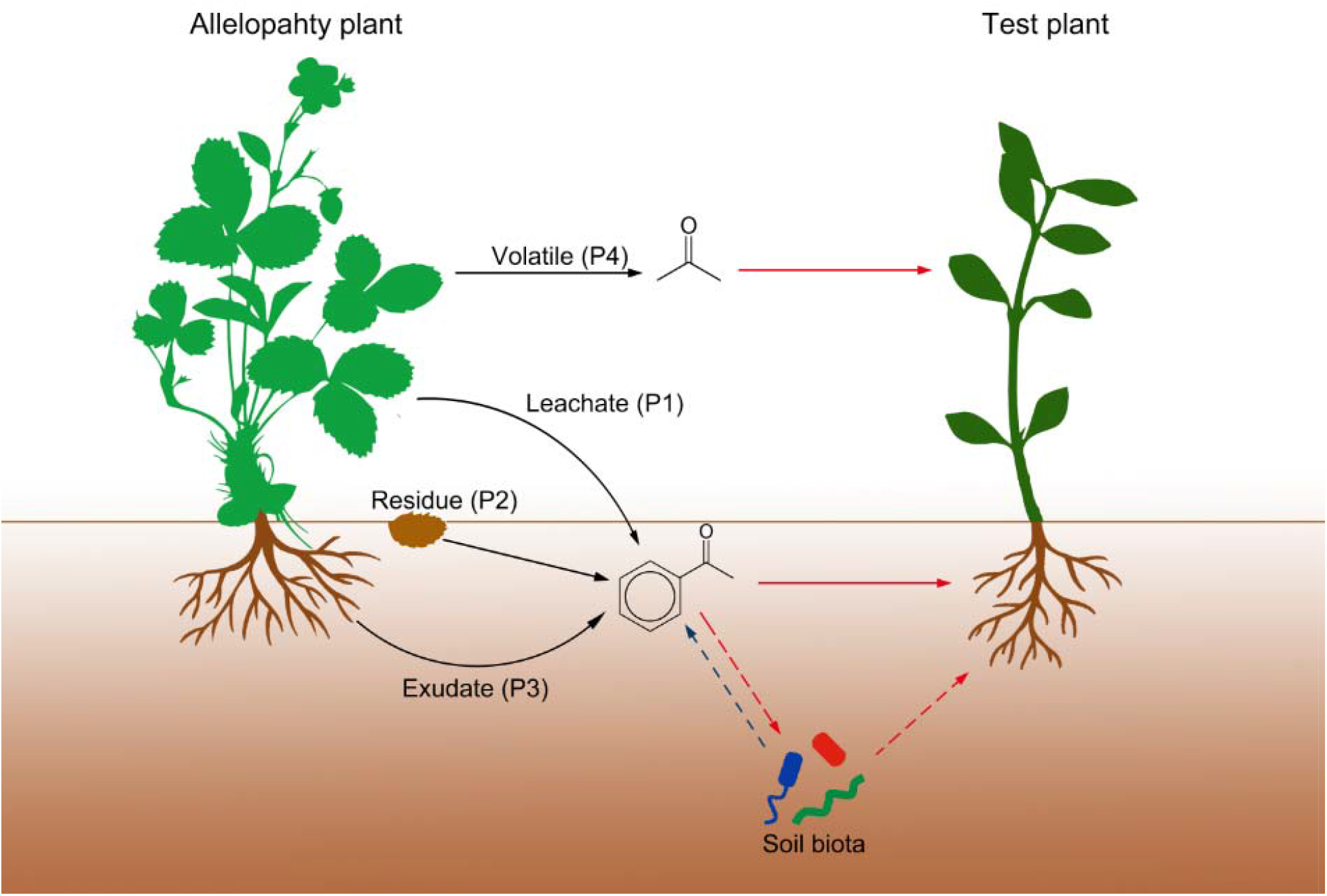
The different release pathways and effects of allelochemicals. The allelopathy plant (left) can release allelochemicals through four pathways (black arrows): leaching by rain (P1), decomposition of plant residues (P2), exudation from roots (P3) and volatilization (P4). The allelochemicals can affect the test plant directly (red arrows) or indirectly through their effect on soil biota (dashed red arrows). Soil biota can also affect allelochemicals, such as through conversion or degradation of allelochemicals. The root illustration is designed by Brgfx/Freepik

**Figure 2.**
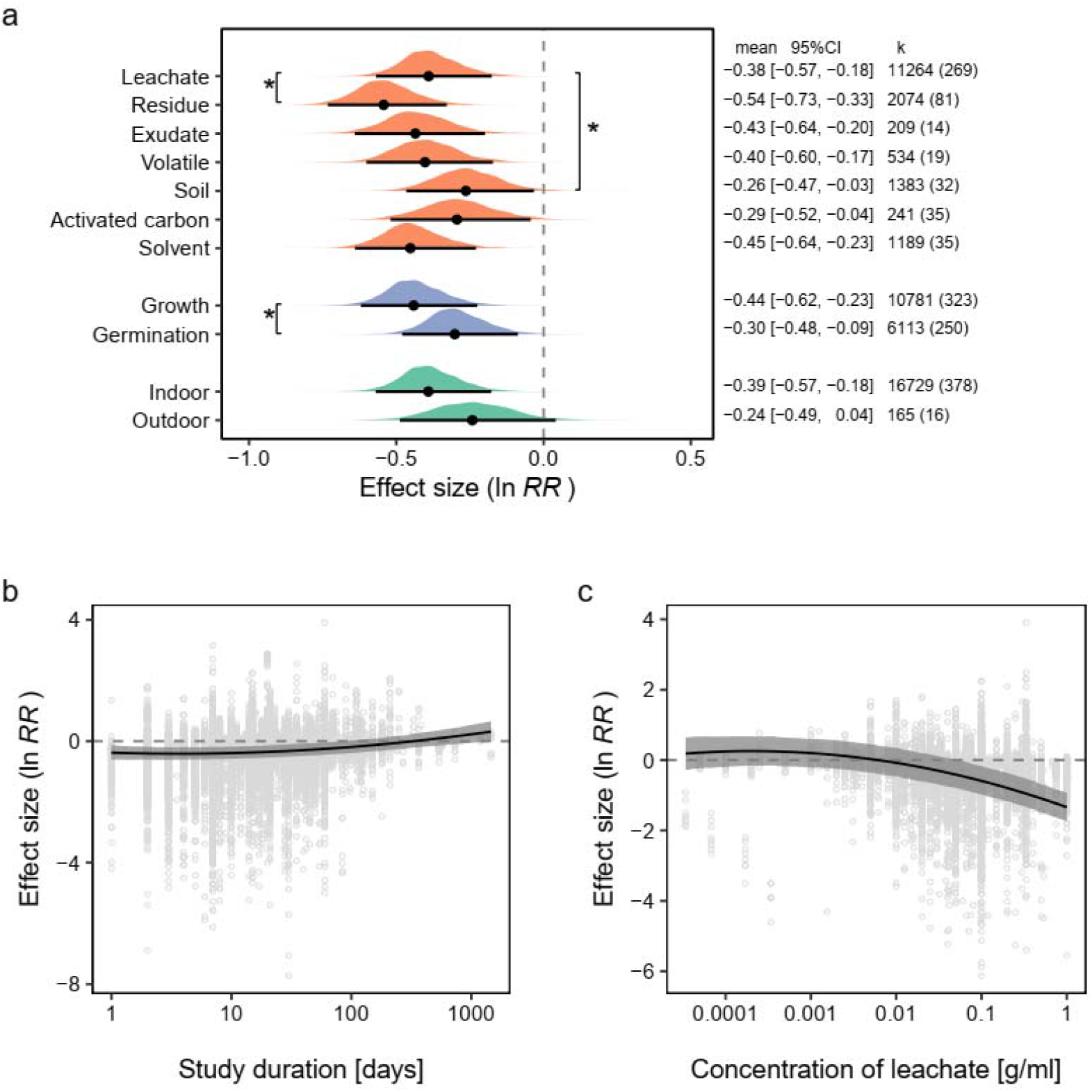
Effects of different aspects of study design on allelopathy. **a**. Effects of the type of method (orange), the type of performance measure (blue) and the experiment environment (green) on allelopathy. **b**. Effect of study duration on allelopathy. **c**. Effect of the concentration of leachates on allelopathy. In **a**, for each category (parameter), the posterior distribution is plotted with the mean and 95% credible interval. The text on the right displays the mean, 95% credible interval (CI), and the number of observations and papers (k). An asterisk indicates a significant difference between the level of interest and the reference level. In **b** and **c**, curves of the estimated effects are shown with their 95% credible intervals. Negative values of the effect size (ln *RR*) indicate that allelopathy inhibits plant performance.

Variance in allelopathy can also arise from biological traits of plants. Several studies of plant succession revealed that the effect of allelopathy may depend on life history. For example, Jackson and Willemsen^17^ reported that ragweed (*Ambrosia artemisiifolia*) and wild radish (*Raphanus raphanistrum*), which are short-lived and early-successional species, cannot reestablish on soils on which the long-lived and late-successional species *Aster pilosus* grew previously. Besides life history, domestication may also matter as crop species appear to be more sensitive to allelopathy than weeds^18^. From an evolutionary perspective, individuals of short-lived species are associated with other species for a shorter period than long-lived species are^19^. Likewise, crop species are less exposed to interference with other species (e.g. weeds) as they are frequently protected by humans. Consequently, the abilities to inhibit other species and to tolerate inhibition from others is less likely to be selected for in short-lived species or crops^16^. However, whether this is true remains untested at a broad scale.

The strength and direction of allelopathic interactions may also depend on two other aspects of evolutionary history: phylogenetic distance and species origin. On the one hand, the composition of secondary metabolites (i.e. allelochemicals) is generally phylogenetically conserved in plants^20,21^. Therefore, distantly related species might strongly inhibit each other due to little overlap in, and thus novelty of, their allelochemical profiles. On the other hand, evolutionary history is also shaped by the conditions that a species has experienced. In their native range, alien plants interacted with other species (e.g. competitors and enemies) than they do in their non-native range^22–24^. Therefore, the aliens might have evolved different arsenals of allelochemicals to which the natives, in the non-native range of the aliens, are not adapted. This is known as the Novel Weapons hypothesis^25^. For example, Callaway and Aschehoug^6^ found that the Eurasian *Centaurea diffusa*, a noxious invasive weed in North America, had more negative effects on species native to North America than on species native to Eurasia. The other way around, the natives may also produce arsenals of allelochemicals to which introduced aliens are not adapted, and this might provide biotic resistance^26–28^. Still, the importance of phylogenetic distance and novelty for the strength of allelopathic interactions and the consequences for invasion success of alien plants remains unknown.

Here, we conducted a meta-analysis to access the overall effect of allelopathy, and identified sources of variance in allelopathy by testing whether allelopathy is related to study design, biological traits of species, and evolutionary history. In addition, we identify some of the knowledge gaps in present allelopathy research, and make suggestions for future direction.

## Results

Our meta-analysis showed that allelopathy was, overall, inhibitory, as it reduced plant performance by on average 25.0% (mean ln RR = −0.288, 95% CI = [−0.492, −0.052]). As expected, the heterogeneity in effect size was high. Within-study heterogeneity (i.e. at the observation level) accounted for 53.6% of the variance, between-study heterogeneity accounted for 26.1% of the variance, between-test-species heteorogeneity (i.e. the non-phylogenetic part) for 13.1% of the variance, and between-allelopathy-species heterogeneity for 7.1% of the variance (Table S1). The phylogenetic signal was low, both among allelopathy species (*H*^*2*^ < 0.01%) and among test species (*H*^*2*^ = 0.02%; Table S1).

### Effects of study design on allelopathy

The type of method significantly affected the effect of allelopathy (Fig. 2a; Table 2). Compared to the leachate method, the effect of allelopathy was more negative (−14.8%) when the study used residues (ln *RR*_*residue* − *leachate*_ = −0.160 [−0.216, −0.106]), and less negative (+13.4%) when the study used soil on which the allelopathy species was grown (ln *RR*_*soil* − *leachate*_ = 0.126 [0.030, 0.219]). The effect of allelopathy in studies that used activated carbon, exudate, volatile or solvent extraction did not deviate significantly from those that used leachates.

**Table 2.**
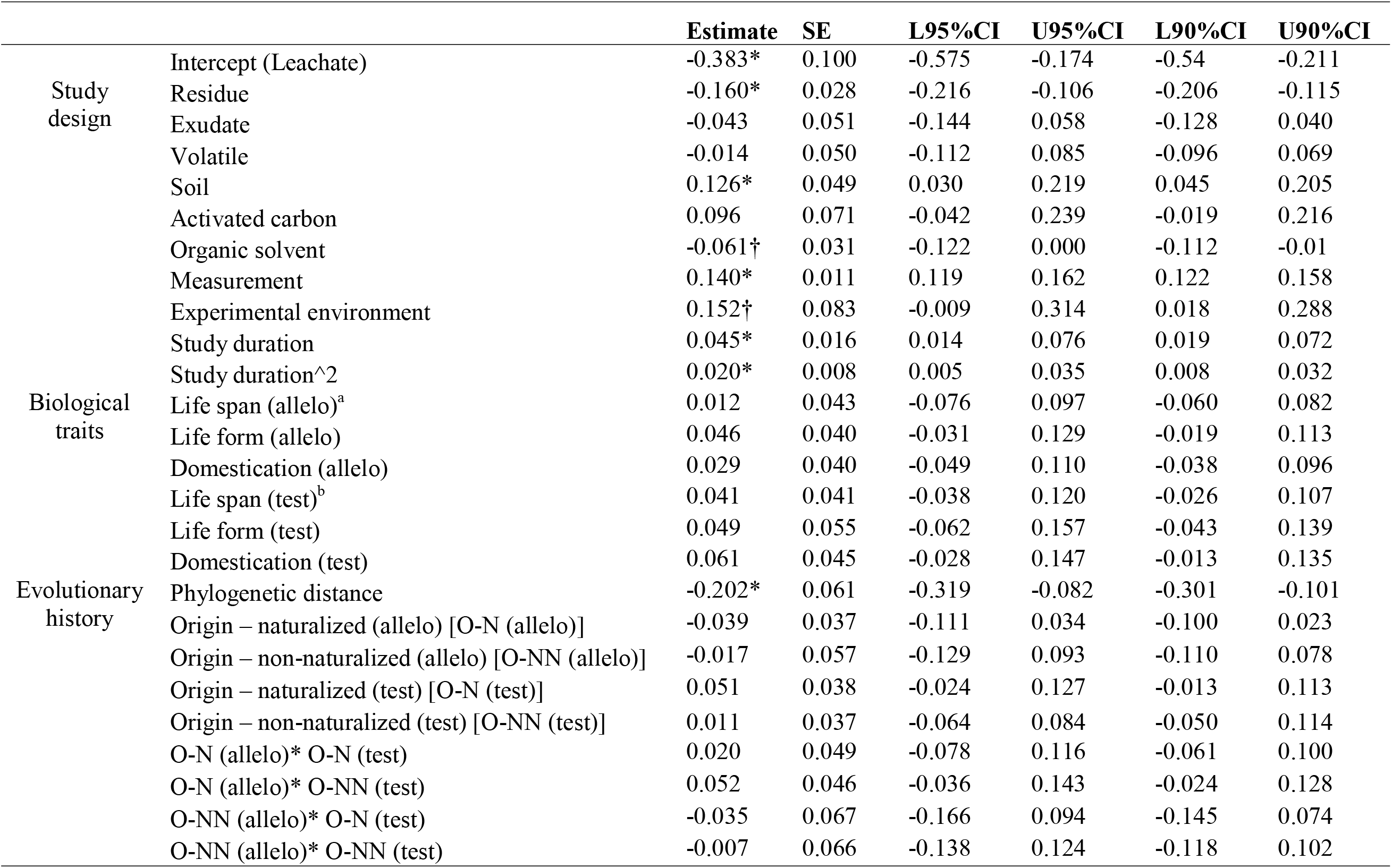
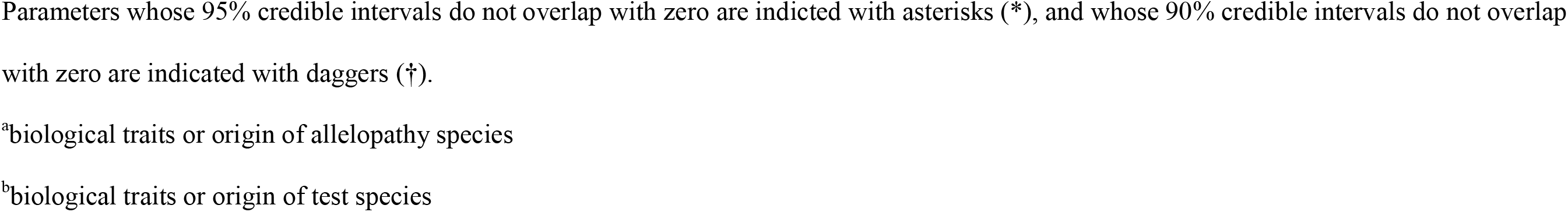
Output of the model testing effects of study design, biological traits and evolutionary history on effect sizes (ln *RR*) of allelopathy. Shown are the model estimates and standard errors as well as the lower (L) and upper (U) values of the 95% and 90% credible intervals (CI).

Allelopathy had overall weaker negative effects (+15.1%) on germination than on plant growth (ln *RR*_*germination* − *growth*_ = 0.140 [0.119, 0.162]; Fig. 2a; Table 2). This was particularly true when allelopathy was tested with leachates (Table S3). However, when tested with residues, allelopathy had stronger negative effects (−25.8 %) on germination than on plant growth (ln *RR*_*germination* − *growth*_ = −0.298 [−0.385, −0.210]; Table S4). Allelopathy tended to have weaker negative effects (+16.4%) in outdoor than in indoor experiments (ln *RR*_*outdoor* − *indoor*_ = 0.152 [−0.009, 0.314]; Fig. 2a; Table 2), but this finding was only marginally significant (90%CI = [0.018 0.288]). We also found that, leachates from belowground parts of plants had weaker negative effects (+13.5%) than leachates from aboveground parts (ln *RR*_*below* − *above*_= 0.127 [0.094, 0.161]; Table S3).

The effects of study duration and concentration of leachates or residues on allelopathy were weakly non-monotonic, as indicated by the significant quadric terms (Fig. 2b-c; Table 2). The effect of allelopathy was negative for experiments of short duration, but its effect was not significantly different from zero when the study duration was over 89 days. The effect of low concentrations of leachates or residues was not significantly different from zero, but the effect became significantly negative when the concentration of leachates exceeded 0.055 g/ml.

### Effects of biological trait on allelopathy

Overall, allelopathy did not differ between short- and long-lived species, between herbaceous and woody species, and between crop and wild species (Fig. 3; Table 2). This holds for both the allelopathy and test species (Fig. 3; Table 2).

**Figure 3.**
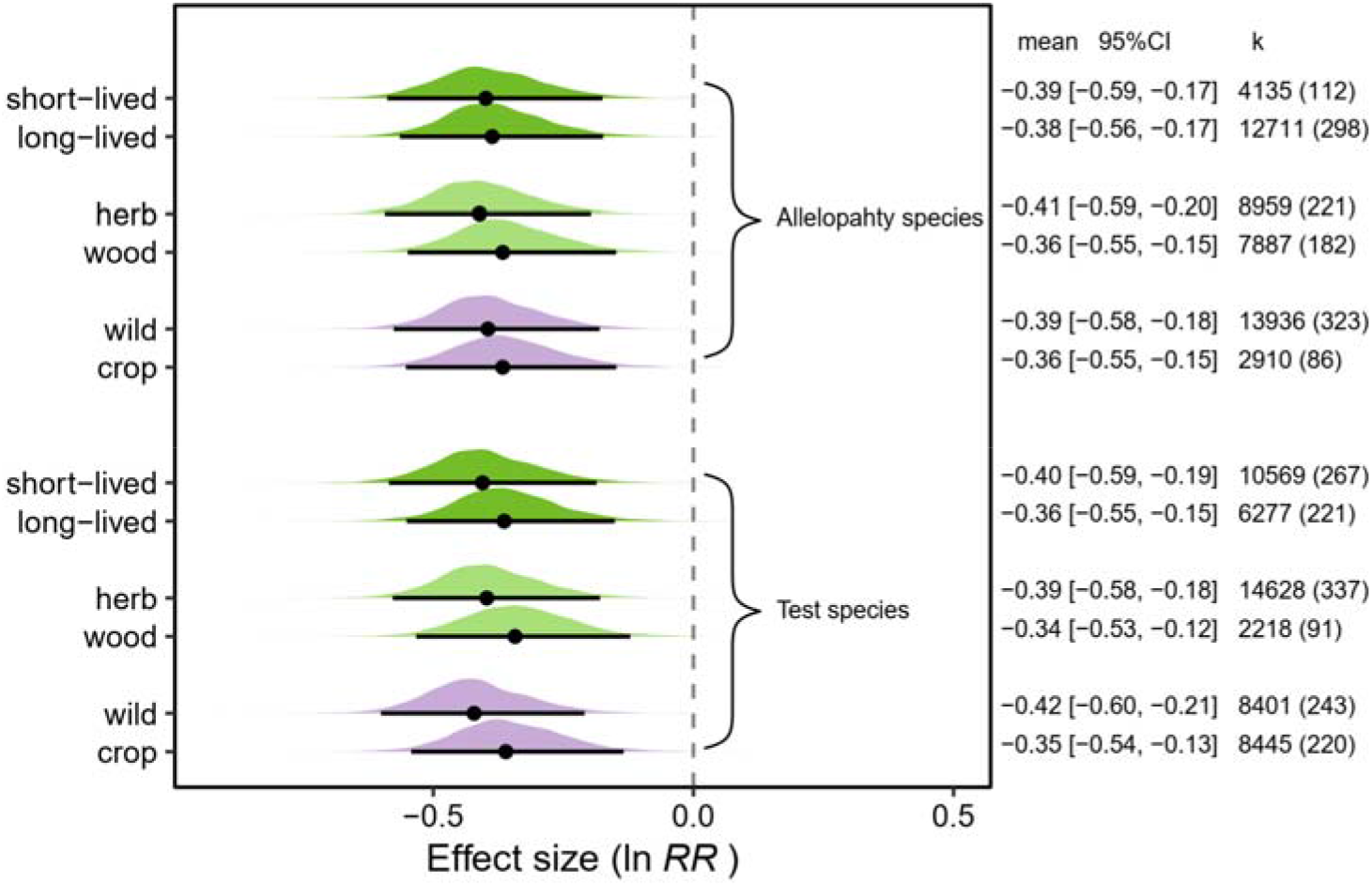
Effects of biological traits (life span, dark green; life form, light green; domestication, violet) on allelopathy. Biological traits of allelopathy species are shown in the upper half, and those of test species in the lower half. For each category (parameter), the posterior distribution is plotted with the mean and 95% credible interval. The text on the right displays the mean, 95% credible interval (CI), and the number of observations and papers (k). Negative values of the effect size (ln *RR*) indicate that allelopathy inhibits plant performance. No significant difference was found.

### Effects of evolutionary history on allelopathy

The effect of allelopathy became slightly more negative as the phylogenetic distance between allelopathy and test species increased (Fig. 4a; Table 2). More specifically, the effect of allelopathy between the most closely related species (or individuals of the same species) reduced plant performance of each other by 27.8% (ln *RR* = −0.327, [−0.513, −0.105]), whereas that between the most distantly related species reduced plant performance by 41.1% (ln RR = −0.529, [−0.730, −0.299]). Across all seven methods (Table 1), allelopathy was not affected by the origin of the allelopathy and test species, or by the interaction between the origins of the two species (Table 2). However, for the subset of studies that tested the effect of leachates, we found that natives experienced more negative (−13.4%) effects from naturalized alien species than from other natives (ln *RR*_*natuarlaized on native* − *native on native*_= = −0.144 [−0.288, −0.002]; Fig. 4b; Table S3). In addition, allelopathic effects between two naturalized aliens tended to be less negative (+13.2%) than allelopathich effects between two natives (ln *RR*_*natuarlaized on naturalized − native on native*_= 0.124 [−0.014, 0.265]; Fig. 4b; Table S3), although this was only marginally significant.

**Figure 4.**
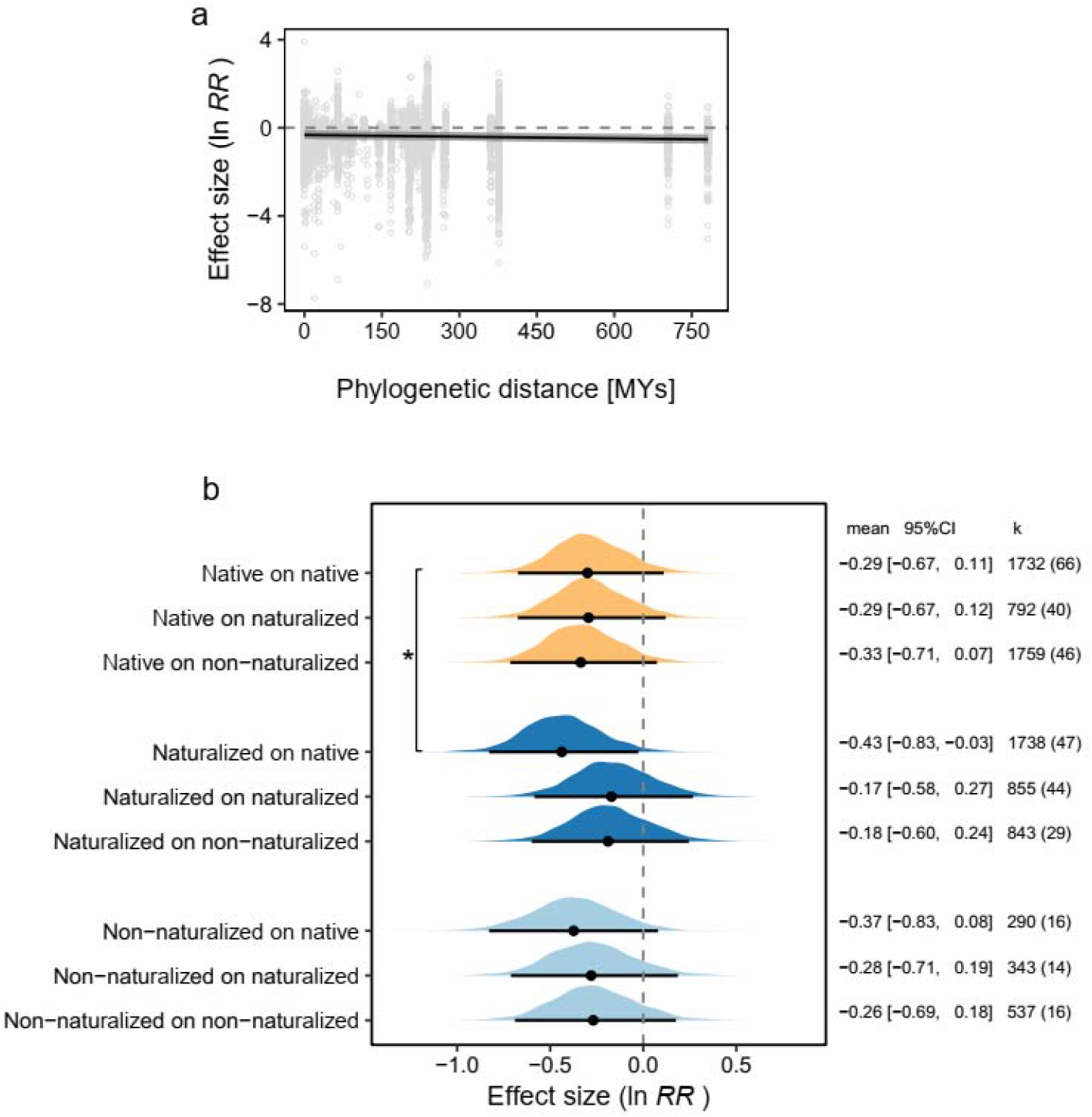
Effects of evolutionary history on allelopathy. **a**. Effect of phylogenetic distance between allelopathy and test species on allelopathy. The curve of the estimated effect is shown with its 95% credible interval. **b**. Effects of origin of allelopathy and test species on allelopathy when tested with the leachate method. Effects of native, non-naturalized alien and naturalized alien allelopathy species are shown in dark blue, light blue and orange. For each category (parameter), the posterior distribution is plotted with the mean and 95% credible interval. The text on the right displays the mean, 95% credible interval (CI), and the number of observations and papers (k). Negative values of the effect size (ln *RR*) indicate that allelopathy inhibits plant performance. An asterisk indicates significant difference between reference level (native on native) and the other.

### Publication bias

The funnel plot of the meta-analysis residuals is asymmetric (Fig. 5a), suggesting some publication bias in the literature. In particular low-precision studies are more likely to report strong negative than strong positive effects, suggesting that the latter are less likely to be published. The presence of a publication bias is confirmed by the Egger’s regression, as the intercept differed significantly from zero (Intercept = −1.466 [−1.678, −1.251]). However, we did not find a significant relationship between effect size and year of publication (Fig. 5b).

**Figure 5.**
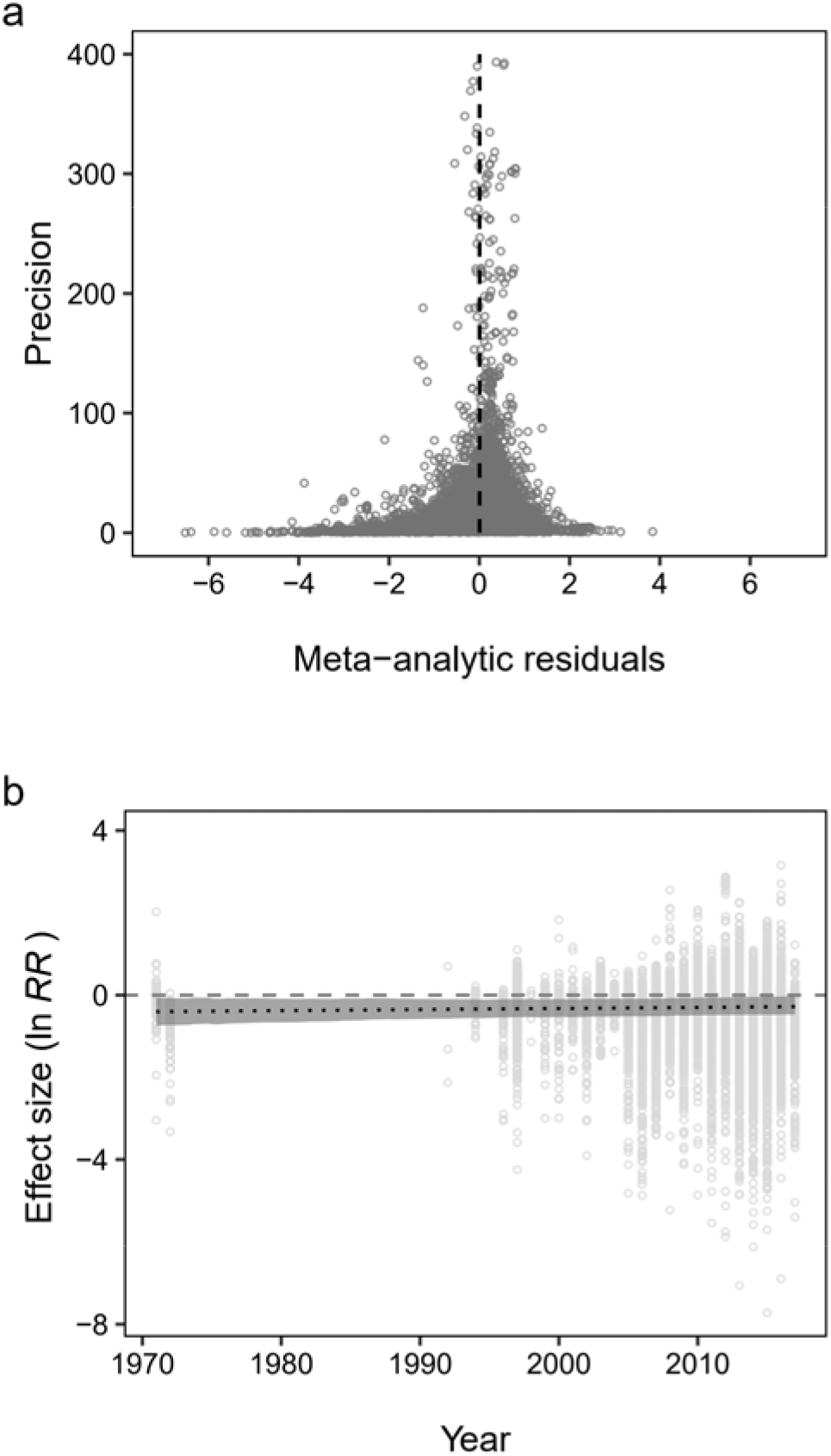
Tests of publication bias for the allelopathy studies used in our meta-analysis. **a**. Funnel plot of the study precision (inverse of standard error) *vs.* the residuals of the meta-analysis. The asymmetry at low levels of precision indicates the presence of a publication of bias. The dashed line was added as visual reference for symmetry. A few data points with extremely high precision are not plotted to aid visualization and a complete plot is provided in the supplement (Fig. S3). **b**. The effect size *vs.* the year of study revealed no significant relationship.

## Discussion

Overall, allelopathy reduced plant performance by 25%. By comparison, competition - which might also include allelopathic effects - reduces plant performance by 43%^29^, and herbivores consume 5% of leaf tissues produced annually by plants^30^. Therefore, allelopathy can be a significant biological force affecting the germination and growth of plants, and consequently the population dynamics of species and community assembly. However, the variation in magnitude and direction of allelopathy is high. Allelopathy was more negative when tested with plant residues than with leachates, and was less negative when tested with soil. Furthermore, the negative effect of allelopathy, generally, became weaker with study duration, and increasingly stronger with the concentration of leachates or residues. Allelopathy was not significantly related to life span, life form and domestication of the two plant species. However, allelopathy became slightly but significantly more negative when phylogenetic distance between allelopathy and test species increased. In addition, native plants suffered more from leachates of naturalized alien plants than from leachates of other native plants, and effects of naturalized alien plants on each other were weak.

### Effects of study design on allelopathy

Allelochemicls can be released into the environment by leaching from plants by rain, decomposition of plant residues, exudation from roots and volatilization (Fig. 2). Our meta-analysis revealed that the effects of exudates and volatiles did not differ from those of leachates, but that plant residues had the most negative effect on plant performance. As the leachates are usually made by soaking plant material in water for *c.* one day, smaller amounts of chemicals might be released than from residues. The leachates treatment is nevertheless not unrealistic, because in nature leaching may only happen during short periods when it rains. Moreover, while leachates contain only water-soluble chemicals, residues could additionally release non-water soluble secondary metabolites, which increases the likelihood that some of those are toxic to test plants. In support of this idea, studies that extracted different fractions of plant secondary metabolites showed that fat-soluble chemicals exerted stronger negative effects than water-soluble ones (Table S9).

Studies that used soils on which allelopathy plants had grown, had weaker negative effects than those that used leachates. Possibly, this indicates the effect of soil microbes on allelopathy^31^. Unlike allelopathy studies that used other methods (Table 1), about 90% of the studies that applied the soil method used soils collected in nature, and thus had soil microbes. A recent study^32^ found that a bacterial species (*Arthrobacter* sp. ZS) isolated from soil greatly increased the degradation of allelochemicals produced by *Ageratina adenophora*. This suggests that soil microbes might neutralize the effect of allelopathy. This could also explain our finding that allelopathy was on average more strongly negative in indoor experiments than in outdoor ones, where soil microbes are probably more diverse and abundant. It is worth noting that few if any studies that used the soil method have ruled out the effect of the allelopathy plant on soil microbes through primary metabolites, which are not involved in allelpathy but could bias the test. For example, consider an allelopathy species that has a negative effect through allelochemicals but fosters, through primary metabolites, arbuscular mycorrhiza fungi. The net effect on the test species might be neutral, and we might wrongly conclude that there was no allelopathy. So, future research needs to account for effects of soil that are not due to allelopathy.

Across the different types of methods, seed germination was less inhibited by allelopathy than plant growth was. This finding is not surprising given that seed germination is mainly determined by seed viability and abiotic environments, including temperature, water and oxygen concentrations^33^, which are unlikely to be affected by the allelopathy treatments. In contrast to the overall pattern, studies that tested allelopathy with plant residues showed that growth was less negatively affected than seed germination. As we also found that plants were less inhibited by litter than by fresh biomass, one potential explanation could be that plant residues (especially litter) also have a fertilization effect that partly compensates the inhibition of plant growth by allelochemicals. Still, effect of allelopathy on fitness remains unclear as fitness was rarely measured.

The negative effect of allelopathy diminished with increasing duration of the experiment, and was absent in most studies that lasted more than 100 days. Possibly, plastic responses of the test plants allow them to compensate for the early negative effects. This could suggest that allelopathy has no long-term impacts. However, if the plants also physically interact, like in nature, the allelopathy species could turn the short-term germination and growth inhibition of its competitors - which gives the allelopathic species more access to resources such as light - into a long-term competitive advantage (e.g. priority effect^34^). As long-term experiments are still rare and allelopathy is rarely tested in combination with other forms of competition (e.g. resource competition, but see studies that used the activated carbon method), this speculation requires further research.

As expected, the inhibitory effect of allelopathy became stronger with increasing concentrations of leachates and residues. An important question then is whether the concentration chosen is realistic. The few studies that give clues^35–39^ show that the annual amount of plant litter varies largely among study sites, ranging from 43 to 884 g/m^2^. Based on this range of concentration, our model predicts that allelopathy through plant residues will reduce plant performance by 2.7% ~ 31.8%, if we assume that the litter ends up in the upper 10 cm of soil and soil bulk density^40^ is 1.5 g/cm^3^. By comparison, our analysis of the residue method showed that allelopathy reduced plant performance by 50% across concentrations (intercept in Table S4). Consequently, the inhibitory effect of allelopathy might be overestimated by some studies that have used unrealistically high concentrations. Towards a more realistic understanding of allelopathic effect, studies that identify the allelochemicals and measure the corresponding concentrations in the soil are necessary.

### Effects of biological trait on allelopathy

Allelopathy was not related to life span, life form or domestication of the species. This was unexpected, because many textbook examples of allelopathy are from long-lived trees, such as *Juglans nigra*^41^ (black walnut) and *Ailanthus altissima*^42^ (tree-of-heaven). Moreover, farmers have long been annoyed by weed interference on crop yield. The reported strong negative effects of some long-lived woody species in nature might come from the accumulation of large amounts of allelochemicals over time, whereas short-lived weedy species leave fewer allelochemicals due to their short generation time and high spatial turnover. When differences in strength of allelopathy between species with different life histories are only due to differences in accumulation of allelochemicals, and not due to differences in toxicity, these differences will not be apparent in short-term experiments with standardized amounts of plant material. On the other hand, the sometimes reported vulnerability of crops might be a result of the crops being grown in monoculture, resulting in a low diversity of allelochemicals, and consequently al lower resistance against invasion by weeds.

### Effects of evolutionary history on allelopathy

Charles Darwin suggested that closely related species are more ecologically similar, and thus will compete more strongly with each other than when they compete with distantly related species (known as Darwin’s naturalization hypothesis^43^). With regard to allelopathy, we found the opposite pattern, i.e. that closely related species (or individuals of the same species) had weaker effects on each other than distantly related species. This finding does not challenge Darwin’s naturalization hypothesis, as the hypothesis is mainly based on resource competition, whereas allelopathy is a form of interference competition. However, the possibly opposing effects of phylogenetic distance on interference and resource competition could help explain why studies testing Darwin’s naturalization hypothesis have not found consistent results^44^.

Still, ecological difference between species, or more specifically the difference in secondary metabolites, cannot be fully captured by phylogenetic distance^45–47^. Secondary metabolites are under selection by competitors and enemies^48,49^, and thus plants that evolved in different regions (i.e. native and alien plants) may not be adapted to each other’s allelochemicals. This might give the alien plants an advantage as posed by the novel-weapons hypothesis^25^ or increase biotic resistance of native communities as posed by the homeland-security hypothesis^26,27^. Although across all types of methods we did not find that the origins of the interacting species mattered, in the studies that used leachates, natives were more strongly inhibited by naturalized aliens than by other natives. On the other hand, the effect of natives on others did not depend on the native-alien status of the other species. So, these findings are in line with the predictions of the novel-weapons hypothesis but not with the predictions of the homeland-security hypothesis.

Surprisingly, although naturalized aliens usually do not share a co-evolutionary history with other naturalized aliens (unless they have the same native range), their leachates had weaker effects on each other than was the case for the effects of natives on other natives (Fig. 4b). The exact reason for this is not clear, but it could imply that naturalized species are less likely to suppress other naturalized species than that they suppress natives. This could contribute to invasional meltdown^50^.

### Future directions

Our meta-analysis of 16,846 effects sizes from 385 studies revealed some clear patterns, but also that there is still lots of unexplained variation in the effects of allelopathy. The explanatory variables regarding study design, and biological traits and evolutionary history of the species accounted for only 3.5% of the variance, and 26.1%, 13.1% and 7.1% of the variance was explained by identity of study, test and allelopathy species, respectively. This means that about 50% of variance remains unexplained. Although this partly reveals the prevalence of heterogeneity in ecology and evolution in general, it might also indicate that some other potentially important explanatory factors were not included. Below we suggest several directions for future research that might help to better understand which factors determine the direction and strength of allelopathy.

#### Positive effects of allelopathy

Molisch^1^ defined allelopathy as the positive or negative effects resulting from biochemical interactions. However, Rice^51^, in the first edition of his influential book ‘Allelopathy’, restricted the definition to harmful effects. Although in the second edition, Rice re-embraced Molisch’s definition, the narrow-sense definition of allelopathy may explain the publication bias in favor of studies that found negative effects. As a result, some of our findings might be biased. For example, if studies that tested low concentrations of allelochemicals found positive effects, they might not have published this result or the publication might not use the term ‘allelopathy’. Actually, in the field of agriculture and horticulture, there is already interest in plant extracts (so-called biostimulants or bioeffectors) that could stimulate crop production^52^. Due to the bias in publication of positive effects, non-monotonic effect of concentration, if present, might be hard to detect using meta-analysis. Therefore, to move towards a more accurate understanding of allelopathy, we should follow Molisch’s original definition of allelopathy, which covers both positive and negative effects.

#### The role of soil microbes

Soil microbes have long been acknowledged to mediate allelopathy^1,2,4^ (Fig. 2). For example, Stinson, et al.^53^ showed that allelochemicals of the non-mycorrhizal European invader *Alliaria petiolata* suppressed growth of native North American plants by disrupting their mutualistic mycorrhizal associations. Microbes might, however, also be in other ways a missing link in studies, especially indoor ones, that test for allelopathic interactions between alien and native species. In a greenhouse experiment, for example, the environment will be novel to both alien and native species, and soil microbes might be absent or foreign to both the alien and native species. Yet, empirical tests that specifically test the effects of soil microbes on allelopathic interactions between plants are still rare. Four out of five direct tests found that the presence of soil microbes had neutralizing effects on allelopathy^14,54–57^. Moreover, our meta-analysis showed that allelopathy was less negative when soil microbes were likely abundant, as in outdoor experiments or studies that used the soil method. However, these results should be interpreted with caution, because our finding provides only indirect evidence, and because three of the direct tests removed soil microbes with autoclaving, which as a side effect can release nutrients^58^. Inoculation with soil microbes or ionizing radiation might be more appropriate methods to manipulate the presence or amount of microbes. In addition, with sequencing techniques becoming more and more affordable, we have more and better tools to study the soil microbiome and its role in allelopathy.

#### Test species

Much of the work on allelopathy has focused primarily on the allelopathy species, as indicated by the fact that fast-growing crops (e.g. lettuce, wheat and rice) have been frequently used as test (i.e. indicator) species. We found, however, that the identity of test species explained nearly twice as much variance in the effects of allelopaty as the identity of allelopathy species did (13.1% *v.s.* 7.1%). Therefore, a stronger focus on the test species may provide new insights. Indeed, how test species respond to allelopathy is important in many circumstances, such as during succession and establishment of alien plants. If a species enters a certain community as a single individual, its establishment will initially, due to its low abundance, hardly depend on how strongly it can affect other species. What matters then is how the individual responds to the communities (e.g. allelopathic effects of the resident plants on the invader). This is a key component of invasion growth rate^59^ (or growth rate when rare).

### Conclusions

Studies on allelopathy have rapidly accumulated since the 1970s. Despite the detection of a potential publication bias, our global synthesis demonstrates that, on average, allelopathy reduced plant performance. Among the four pathways through which allelochemicals are released into the environment, plant residues exerted the most negative effect. Still, the importance of allelopathy in nature requires further investigation, as effects of allelopathy were weaker in studies where soil microbes were likely to be abundant, and when study duration was longer. Finally, we found that closely related species (or individuals of the same species) had weaker allelopathic effects on each other than distantly related species. This suggests that allelopathy would favor coexistence of closely related species (or dominance of single species), which is opposite to resource competition. Therefore, investigating allelopathy and resource competition simultaneously will advance our understanding of species coexistence.

## Methods

### Literature search

We searched *ISI Web of Science* for between-plant-allelopathy studies that were published between January 1900 and December 2017. We used the search string ‘allelopath* AND plant’, which resulted in 3,699 papers. This list of papers was further expanded by including all papers that were published in the *Allelopathy Journal* (ISSN 0971-4693) between 1994 (the year that the journal was first published) and 2017, which resulted in another *c.* 900 papers. We then screened the titles, abstracts and main texts. Studies were included if the following criteria were met:

1. The study tested a pairwise interaction, that is, the effect of one species on another or on conspecifics. Therefore, studies that tested effects of allelopathy on a community, that tested effect of mixed species, and that tested effects of single allelochemicals on plants were excluded.
2. The study tested allelopathy using one of the seven major methods listed in Table 1.
3. The study measured traits related to germination, growth or fitness, and thus studies that measured other traits, such as mycorrhizal infection and the concentration of chlorophyll, were excluded. Germination measurements include germination rate (i.e. proportion of seeds that germinated), germination speed and time to germination. Growth measurements include total biomass and biomass of different plant parts (shoot, roots, leaves and stem); length of roots and shoot; numbers of leaves and branches; leaf size; and stem diameter. Fitness measurements include biomass of seeds, fruits and flowers, and numbers of seeds, fruits, flowers and ramets.
4. The study used seed plants. The few studies that used algae or ferns were excluded.
5. The study reported the duration of the experiment.
6. Mean values and variances (standard deviations, standard errors, or 95% confidence intervals) of control and allelopathy treatments can be obtained from the text, tables, figures or supplements.

### Data collection

In total, 379 published papers met our criteria. We also added data of six experiments from our labs, two of which were published after 2017^60,61^. So, we included 385 studies in total. If a study sampled data at multiple time points, we only extracted the final data points to avoid temporal autocorrelation. If a study tested multiple species pairs, applied multiple treatments (e.g. different concentration of leachates), we considered them as independent observations. For each observation, we extracted the mean values, variances, and sample sizes of the test species, under control and allelopathy treatments. Data from figures were extracted using ImageJ^62^. In addition, we recorded the experiment environment (outdoor and indoor), the plant part used for making the allelopathy treatment (shoot, root and whole plant), study duration, and the concentration of leachates or residues if it was reported in the paper. Our final data set contained 16,846 data points covering 461 allelopathy species and 470 test species.

In addition to the data extracted from the papers, we acquired three biological traits - life span (short- or long-lived), life form (woody or non-woody) and domestication (crop or wild) - of the species by matching their names with several databases. Before matching our database with others, we harmonized taxonomic names of all species according to The Plant List (http://www.theplantlist.org/), using the *Taxonstand* R package^63^. We extracted the information on life span and life form from the following databases: the TRY database^64^(last accessed on 9 November 2015), World Checklist of Selected Plant Families (WCSP, http://wcsp.science.kew.org/, last accessed on 22 August 2018), Plants for a Future (PFAF, https://pfaf.org/, last accessed on 22 August 2018), the LEDA^65^(last accessed on 22 August 2018) and the PLANTS database (http://plants.usda.gov/, last accessed on 22 August 2018). We classified annual and biennial species as short-lived, and perennials as long-lived. A few species were not included in the databases (29% in case of life span and 16% in case of life form). For them, we extracted the information on life span and life form from the description of the species in the allelopathy papers or we did internet searches to find this information. We classified a species as a crop species if it is recorded as a human food in the World Economic Plants (WEP) database (National Plant Germplasm System GRIN-GLOBAL; https://npgsweb.ars-grin.gov/gringlobal/taxon/taxonomysearcheco.aspx, last accessed on 7 January 2016), which is based on a book by Wiersema and León^66^.

We also acquired data on the origin of the species (i.e. native or alien at the location of study), and in the case of alien species, we distinguished between non-naturalized and naturalized species. The latter are species that have established self-perpetuating populations in the wild at the location of study (sensu Richardson et al.^67^). For this, we used three databases – the Global Naturalized Alien Flora (GloNAF), the Germplasm Resources Information Network (GRIN), and the Plants of the World Online portal (POWO). The GloNAF database contains lists with naturalization status of 13,939 alien vascular plant taxa for 1,029 regions (countries or subnational administrative units)^68^. GRIN is an online database published by the United States Department of Agriculture (USDA, https://www.ars-grin.gov/, last accessed on 2 March 2020). POWO is an online database published by the Royal Botanic Gardens, Kew (http://powo.science.kew.org/, last accessed on 2 March 2020). All of them provide descriptions of the distribution of the taxa. We classified a species at the location of study as a naturalized alien if GloNAF or GRIN reported that it is naturalized there; as native if GRIN, POWO or the study reported that it is native there; as non-naturalized alien if the species is recorded there neither as native nor as naturalized by any of the data sources. Five studies tested for allopathic effects of an alien species collected in its non-native region on species that had been collected in the native region of the alien species^6,60,69–71^. As in these cases, it is not clear whether the allelopathy and test species should be considered native or alien, we excluded such species pairs from analysis.

We constructed a phylogenetic tree for all species in our dataset (Supplement S5) by pruning an existing phylogenetic supertree using the R function *S.PhyloMaker*^72^. The supertree, which was initially constructed by Zanne, et al.^73^, and further corrected and expanded by Qian and Jin^72^, included 30,771 seed plants, and has been time-calibrated for all branches using seven gene regions (i.e., 18S rDNA, 26S rDNA, ITS, *matK*, *rbcL*, *atpB*, and *trnL-F*) available in GenBank and fossil data. Species that were absent in the supertree (about 21.6%), we added as polytomies to the root of their genus or family. We calculated the pairwise phylogenetic distance between allelopathy and test species using the *cophenetic* function in the *ape* package^74^.

### Effect size calculation

As effect size, we calculated the log response ratio (ln *RR*)^75^ as:

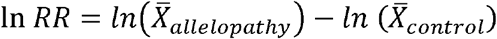

where 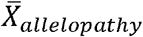 and 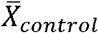 are the mean values of performance (germination, growth or fitness) of a test species in the allelopathy and control, respectively. When a high value of a measured trait indicates low performance of plants (e.g. For a given species, a high value for germination time indicates low success of germination), we changed the sign of the effect sizes of this trait. Consequently, a negative value of ln *RR* indicates that allelopathy reduces plant performance, and a positive value indicates that allelopathy increases plant performance.

The sampling error variance of the ln *RR* was calculated as:

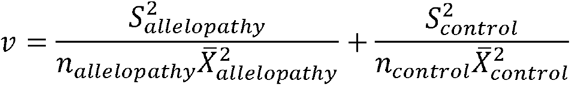

where *S* and *n* are the standard deviation and sample size, respectively^75^.

### Statistical analyses

All analyses were carried out with Bayesian multivariate models implemented in the *brms* package^76^ in R 3.6.1^77^, which allowed us to account for phylogenetic non-independence of the species and to include both the sampling error and the residual. We considered an estimated parameter as significant when its 95% credible interval of the posterior distribution did not overlap zero, and as marginally significant when the 90% credible interval did not overlap zero.

### Meta-analysis

To test the overall effect of allelopathy, we ran an intercept-only meta-analysis model. We included effect size (ln *RR*) as response variable, and study identity (i.e. paper), identities and phylogenetic effects (added as phylogenetic variance-covariance matrices) of allelopathy and test species, and sampling error as random effects. Note that both the identity and phylogenetic effect of species are species-specific effects, but the former represents the non-phylogenetic part^78^.

We used the default priors set by the *brms* package, but changing the priors had negligible effect on our results. We ran four independent chains with 3,000 iterations each. Half of the iterations were used as warm-up, so a total of 6,000 samples were retained to estimate the posterior distribution of each parameter. We quantified heterogeneity of the model by estimating *I*^*2*^ *sensu* Nakagawa and Santos^78^ (see Supplement S2 for details). The phylogenetic signal was quantified by *H*^*2*^, which is a metric of phylogenetic signal named phylogenetic heritability by Lynch^79^. Similar to the widely used metric of phylogenetic signal Pagel’s λ^80^, *H*^*2*^ = 0 indicates that there is no effect of phylogenetic relatedness of species on their difference in effect sizes, while *H*^*2*^ = 1 indicates that the differences in effect-sizes are exactly proportional to the phylogenetic relatedness of the species. To report the effect of allelopathy on plant performance at the raw-data scale, we back-transformed the log response ratio (ln *RR*) with the formula: e^ln *RR*^ − 1. For example, ln *RR* = −0.5 indicates that allelopathy reduced plant performance by 39.3% (e^−0.5^ − 1).

### Meta-regressions

To test the importance of different sources of variance in allelopathy, we ran three meta-regression models (i.e. mixed-effects meta-analyses). The first model included data from all seven methods described in Table 1. The second and third models analyzed two subsets, that is, the leachate and the residue method, respectively, which allowed us to analyze certain aspects of the experimental design in more detail. We also ran meta-regression models for each of the other five methods, but because the number of studies was relatively small, we report their results in the supplements (Supplement S3). In all models, we included effect size as response variable; and used the same random effects, priors, warm-up, and sampling in the intercept-only meta-analysis described in the ‘Meta-analysis’ subsection above.

In the first model, we included the following explanatory variables as fixed effects: **1)** type of method (leachate, residue, activated carbon, solvent, soil, volatile and exudate; see Table 1); **2)** type of performance measure (germination and growth; as only 10 studies measured fitness, we combined fitness with growth); **3)** experiment environment (indoor and outdoor); **4)** study duration (days); **5)** life span (short-lived [annual and biennial] and long-lived [perennial]) of allelopathy and test species; **6)** life form (herbaceous and woody) of allelopathy and test species; domestication (crop and wild) of allelopathy and test species; **8)** phylogenetic distance between allelopathy and test species; and **9)** origin (native, naturalized alien and non-naturalized alien) of allelopathy and test species, and interaction between origin of allelopathy and test species.

Variables 1-4 are those related to the experimental design. Variables 5-7 are those related to biological traits. Variables 8 and 9 are those related to evolutionary history. Note that the variables 5-7 (life span, life form and domestication) are also related to evolution. For example, crops were selected for high yield by humans. However, the variables 8 and 9 (phylogenetic distance and origin) focus on the similarity in evolutionary history between allelopathy and test species, for example, the relative position on the phylogenetic tree between two species.

In the second and third models, which focused on studies that used the leachate and residue methods, respectively, we included the same fixed effects as in the first model, except for type of method. Consequently, the models included the following fixed effects: **1)** type of measurement; **2)** experiment environment (not for leachate, because there was no outdoor experiment); **3)** study duration; **4)** life span of allelopathy and test species; **5)** life form of allelopathy and test species; **6)** domestication of allelopathy and test species; **7)** phylogenetic distance between allelopathy and test species; and **8)** origin of allelopathy and test species, and their interaction.

We included another three fixed effects related to the study design, which could not be included in the first meta-regression model: **9)** ‘color’ of biomass (fresh biomass [green] and litter [brown]); **10)** plant part (belowground, aboveground part and whole plant); **11)** concentration of leachates or residues.

Because the concentration of secondary metabolites, study duration and phylogenetic distance have been found and/or hypothesized to be nonlinear^2,81,82^, we considered nonlinear effect for all of them. To do this, we first ran models that applied data transformation (natural-log and square-root transformation), included quadric terms or specified non-linear curves (Monod equation and Weibull curve). Then, we performed model selection with the leave-one-out cross-validation method using the *loo* package^83^. In the end, we natural-log transformed study duration and concentration of leachates or residues and added quadric terms for those variables, while we used the original linear scale for phylogenetic distance. To improve comparability of the coefficient estimates, we bounded phylogenetic distance, which ranges from 0 (i.e. same species) to 781 million years, between 0 and 1.

To improve interpretation of the intercept^84^ and to aid comparisons between different meta-regression models, we mean-centered all categorical explanatory variables with two levels after coding them as dichotomous (0 and 1) variables. For categorical variables with more than two levels, we set the most common levels as the reference levels. For type of method this was ‘leachate’, for plant part this was ‘aboveground part’, and for origin of allelopathy and test species this was ‘native’. For all continuous variables, we applied median-centering instead of mean-centering because their distributions were skewed.

For the concentration of leachates or residues, studies used three types of units (weight/weight, g/g; weight/volume, g/cm^3^; and weight/area, g/cm^2^), which differ in their scales and might have different per-unit effects. To account for this, we included type of unit and set ‘weight/weight’, the most common level, as the reference level in the model for residue method. In the model for leachate method, all but two studies used the unit of weight/volume, and we therefore removed the two exceptions from the analysis. Finally, of the studies that used the residue method, only a few used non-naturalized alien species. For example, only two studies tested effect of non-naturalized aliens on natives. Therefore, we did not discriminate between non-naturalized and naturalized aliens in the model for residue method.

### Detection of Publication bias

Because statistically significant results are more likely to be published than non-significant results^85^, publication bias can affect the results of meta-analyses. To test for publication bias, we first used the funnel-plot method by plotting meta-analytic residuals against the inverse of their precision (i.e. inverse of the sampling error). When the funnel plot is asymmetric, one can conclude that publication bias is present. The meta-analytic residual is a sum of within-study effect (analogy to residual in linear models) and sampling-error effects^78^, and was extracted from the meta-analysis (the intercept-only model). Second, we performed a modified Egger’s regression^78,86^ by using the meta-analytic residuals as the response variable, and the precision as the explanatory variable. When the intercept does not overlap zero, one can conclude that publication bias is present. Finally, we analyzed the temporal trends in effect sizes. A so-called time-lag bias arises when the earliest published studies have larger effects than those of later studies^87^. To do so, we ran a meta-regression model that included effect size as response variable, and publication year as the only fixed effect.

## Supporting information

all supplements

## Acknowledgements

We thank V. Pasqualetto for help with data extraction; and R. Hoefer, J. Kern, Y. Ling, O. Michels, A. Oduor and D. Prati for sharing their data. ZZ acknowledges funding from the China Scholarship Council (201606100049) and support from the International Max Planck Research School for Organismal Biology. YL acknowledges funding from Chinese Academy of Sciences (Y9H1011001, Y9B7041001).

## Author contributions

MvK and EW conceived the idea. ZZ, YL and MvK designed the study. ZZ, YL and LY collected the data. ZZ led the analyzing and writing, with input from all others.

