## Supplementary material for "Effect of allelopathy on plant performance": all supplements

Supporting information

### Supplement S1 Overview of published allelopathy studies.

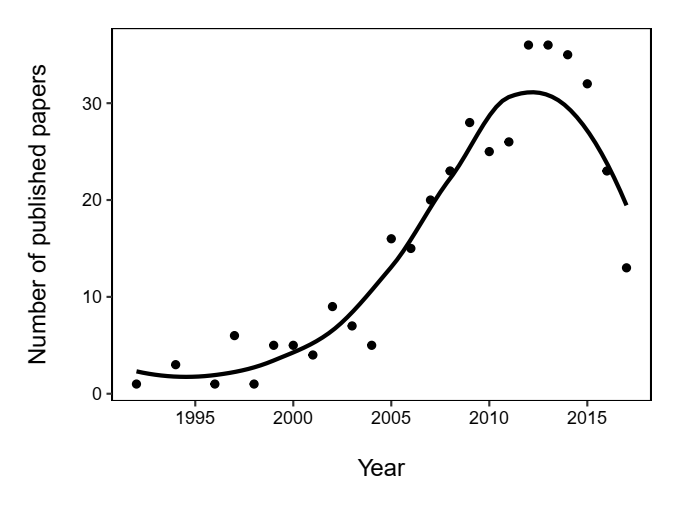

**Figure S1** Temporal trends in studies of allelopathy. Only studies that were included in our final dataset are shown. A LOESS curve has been added to aid visual interpretation.

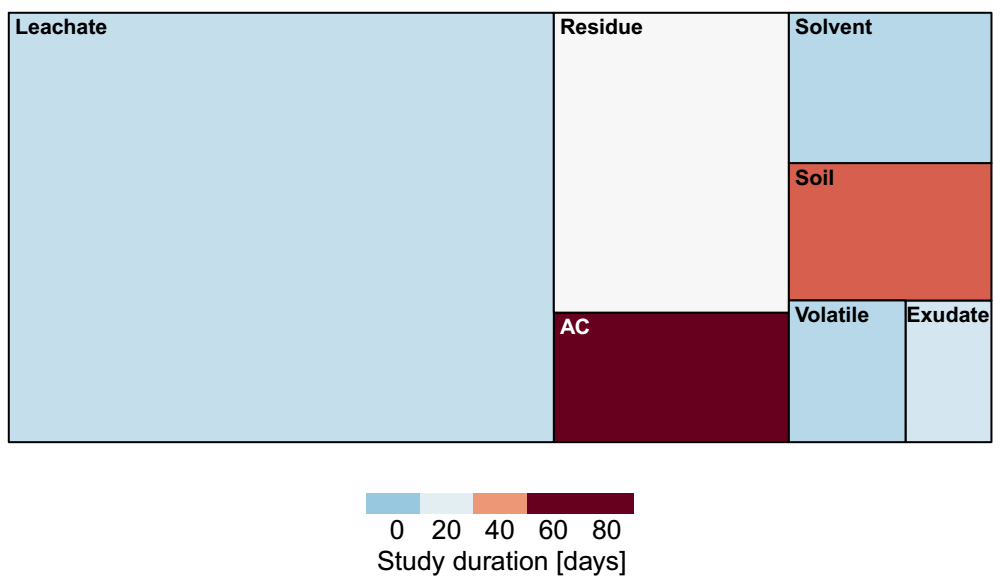

**Figure S2** A treemap showing the number of studies and their median study duration for each type of method (see Table 1) used to study the effects of allelopathy. The size of the rectangle is proportional to the number of studies, and the color of the rectangle indicates the median study duration (days) for each method.

### Supplement S2 Heterogeneity

#### The model

The model of our meta-analysis can be written as:

$$z_{i}= u_{i}+p_{j\left[ i \right]}+ a_{A_{k\left[ i \right]}}+ a_{T_{l\left[ i \right]}}+s_{A_{k\left[ i \right]}}+ s_{T_{l\left[ i \right]}}+ m_{i}+ e_{i}$$

$$\mathbf{p} \sim N\left( \mathbf{0}, \sigma_{p}^{2}\boldsymbol{I} \right)$$

$$\boldsymbol{a}_{\boldsymbol{A}} \sim N\left( \mathbf{0}, \sigma_{a_{A}}^{2}\boldsymbol{A}_{\boldsymbol{A}} \right)$$

$$\boldsymbol{a}_{\boldsymbol{T}} \sim N\left( \mathbf{0}, \sigma_{a_{T}}^{2}\boldsymbol{A}_{\boldsymbol{T}} \right)$$

$$\mathbf{s}_{\boldsymbol{A}} \sim N\left( \mathbf{0}, \sigma_{s_{A}}^{2}\boldsymbol{I} \right)$$

$$\mathbf{s}_{\boldsymbol{T}} \sim N\left( \mathbf{0}, \sigma_{s_{T}}^{2}\boldsymbol{I} \right)$$

$$\mathbf{m} \sim N\left( \mathbf{0},\mathbf{M} \right)$$

$$\mathbf{e} \sim N\left( \mathbf{0}, \sigma_{e}^{2}\boldsymbol{I} \right)$$

where $z_{i}$ is the effect size of observation *i*, and $u_{i}$ is its estimated effect size according to explanatory variables (e.g. when there is no explanatory variable, it is the intercept). $p_{j\left[ i \right]}$ is the study specific effect of study *j*.$a_{A_{k\left[ i \right]}}$ and $a_{T_{l\left[ i \right]}}$ are the phylogenetic effects of allelopathy species *k* and test species *j*, respectively. $s_{A_{k\left[ i \right]}}$ and $s_{T_{l\left[ i \right]}}$ are the specific effect of allelopathy species *k* and test species *j*, respectively. Note that these two represent the non-phylogenetic part. $m_{i}$ is the sampling error of the observation *i*. $e_{i}$ is the residual. The sum of $m_{i}$ and $e_{i}$ is the meta-analytic residual.

$\sigma_{p}^{2}$ is the between-study variance, which is estimated from the data. $\sigma_{a_{A}}^{2}$ and $\sigma_{a_{T}}^{2}$ are the phylogenetic variance of allelopathy and test species, respectively. $\boldsymbol{A}_{\boldsymbol{A}}$ is a N_allelopathy_species_ by N_allelopathy_species_ phylogenetic correlation matrix for allelopathy species, and $\boldsymbol{A}_{\boldsymbol{T}}$ is a N_test_species_ by N_test_species_ phylogenetic correlation matrix for test species. $\sigma_{s_{A}}^{2}$ and $\sigma_{s_{T}}^{2}$ are the between-species variance for allelopathy and test species, respectively. **M** is a N_observation_ by N_observation_ matrix with diagonal being the sampling error variance of each observation. $\sigma_{e}^{2}$ is the between-observation variance (or within-study variance).

#### Calculation of heterogeneity and phylogenetic signal.

Total heterogeneity ($\sigma_{t}^{2}$) is calculated as:

$$\sigma_{t}^{2}= \sigma_{p}^{2}+ \sigma_{a_{A}}^{2}+\sigma_{a_{T}}^{2}+ \sigma_{s_{A}}^{2}+ \sigma_{s_{T}}^{2}+\sigma_{m}^{2}+ \sigma_{e}^{2}$$

where

$$\sigma_{m}^{2}= \sum w_{i} \left( n-1 \right)/\left[ \left( \sum w_{i} \right)^{2}- \sum w_{i}^{2} \right]$$

*w* is the inverse of sampling error variance and *n* is the number of observations. All other symbols are as above.

Then, the amount of heterogeneity at study ($I_{p}^{2}$), allelopathy species ($I_{s_{A}}^{2}$), test species ($I_{s_{T}}^{2}$) and within-study ($I_{e}^{2}$) levels are calculated as:

$$I_{p}^{2}= \sigma_{p}^{2}/\sigma_{t}^{2}$$

$$I_{s_{A}}^{2}= \sigma_{s_{A}}^{2}/\sigma_{t}^{2}$$

$$I_{s_{T}}^{2}= \sigma_{s_{T}}^{2}/\sigma_{t}^{2}$$

$$I_{e}^{2}= \sigma_{e}^{2}/\sigma_{t}^{2}$$

and phylogenetic signal of allelopathy species ($H_{a_{A}}^{2}$) and test species ($H_{s_{A}}^{2}$) are calculated as:

$$H_{a_{A}}^{2}= \sigma_{a_{A}}^{2}/(\sigma_{t}^{2}-\sigma_{m}^{2})$$

$$H_{a_{T}}^{2}= \sigma_{a_{T}}^{2}/(\sigma_{t}^{2}-\sigma_{m}^{2})$$

**Table S1** Amount of heterogeneity and phylogenetic signal of the intercept-only meta-analysis.

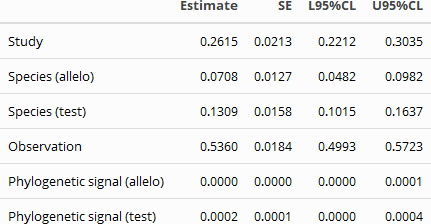

### Supplement S3 Sources of variance in allelopathy for each method

We ran meta-regression models for each of the seven methods that were used to test allelopathy (Table 1). The model for leachate and residue method were described in the main text. The other five models are similar. The explanatory variables included in each of the five model are shown in Table S2, and the results for each of the seven models are shown in Tables S3-S9.

**Table S2** Explanatory variables for each model are indicated by ‘Y’.

|  | Variables | Exudate | Volatile | Soil | Activated carbon | Solvent extraction |
| --- | --- | --- | --- | --- | --- | --- |
| Study design | Measurement | Y | Y | Y | Y | Y |
|  | Experiment environment |  |  |  | Y |  |
|  | Study duration | Y | Y | Y | Y | Y |
|  | Concentration |  |  |  |  | Y |
|  | Solvent |  |  |  |  | Y |
| Biological traits | Life span (allelo) | Y | Y | Y | Y |  |
|  | Life form (allelo) | Y | Y | Y | Y |  |
|  | Domestication (allelo) | Y | Y | Y | Y |  |
|  | Life span (test) | Y | Y | Y | Y |  |
|  | Life form (test) | Y | Y |  | Y |  |
|  | Domestication (test) | Y | Y | Y | Y |  |
| Evolutionary history | Phylogenetic distance | Y | Y | Y | Y | Y |
|  | Origin | Y | Y | Y | Y | Y |

^a^In the model for soil method, life form of test species was not included due to low number of studies that tested woody species.

^b^In the model for solvent-extraction method, biological traits were not included due to their collinearity with study duration and origin of species. The model also included ‘type of solvent’, as the extraction were prepared with different types of solvents.

**Table S3** Effects of study design, biological traits and evolutionary history on effect sizes (ln *RR*) of allelopathy in **leachate** method. Shown are the model estimates and standard errors as well as the lower (L) and upper (U) values of the 95% and 90% credible intervals (CI).

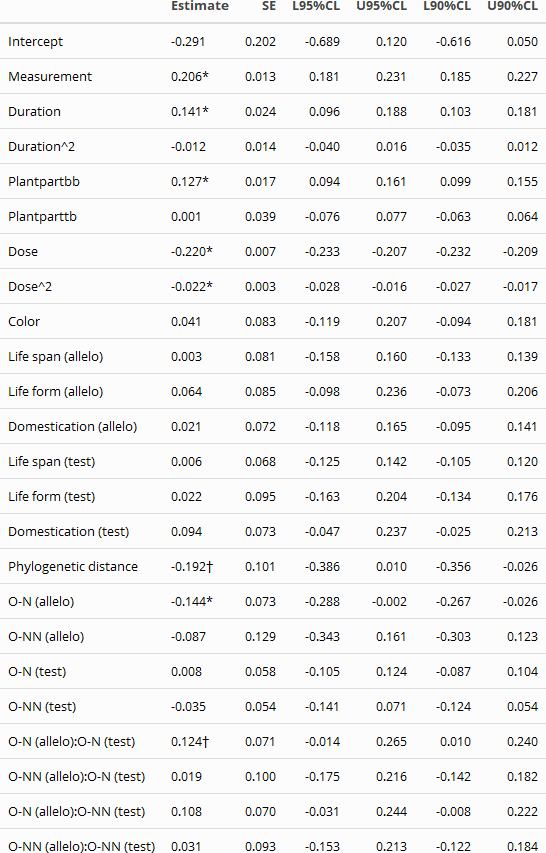

^A^plantpart: bb, belowground biomass; tb, total biomass.

**Table S4** Effects of study design, biological traits and evolutionary history on effect sizes (ln *RR*) of allelopathy in **residue** method.

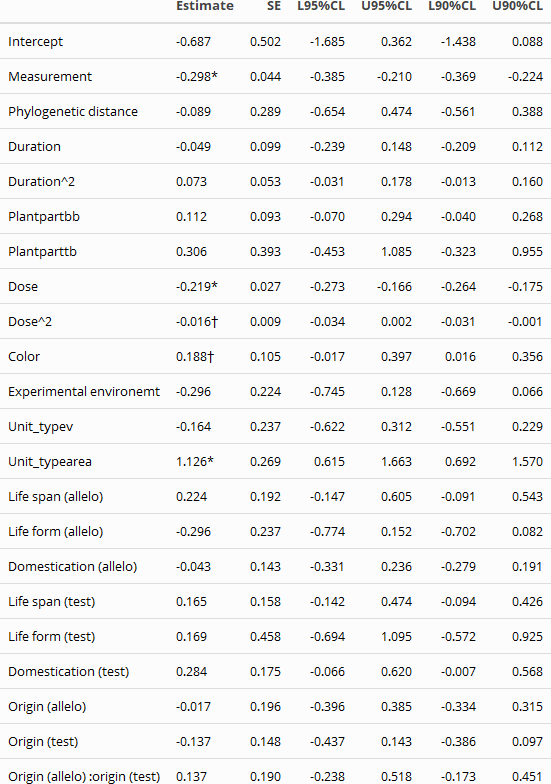

**Table S5** Effects of study design, biological traits and evolutionary history on effect sizes (ln RR) of allelopathy in **exudate** method.

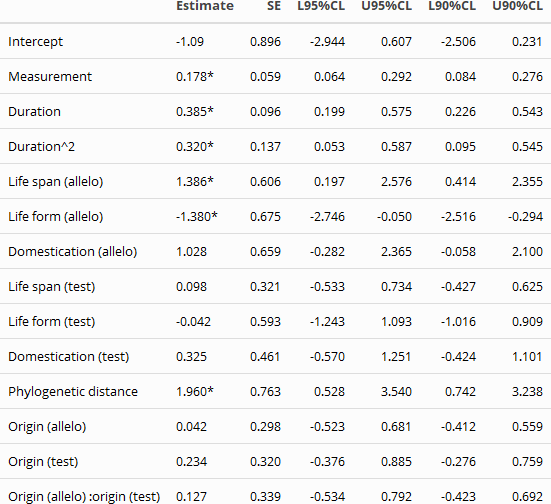

**Table S6** Effects of study design, biological traits and evolutionary history on effect sizes (ln RR) of allelopathy in **volatile** method.

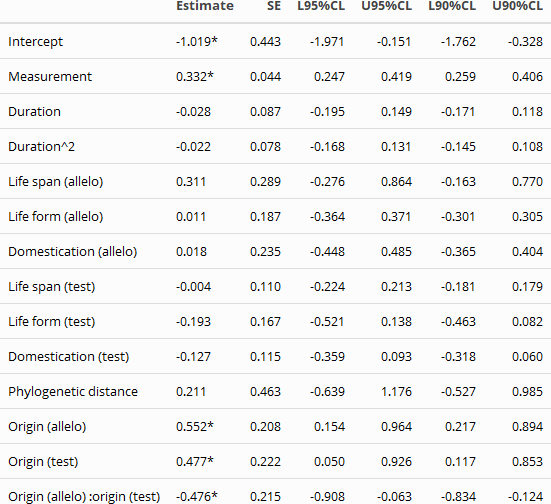

**Table S7** Effects of study design, biological traits and evolutionary history on effect sizes (ln RR) of allelopathy in **soil** method.

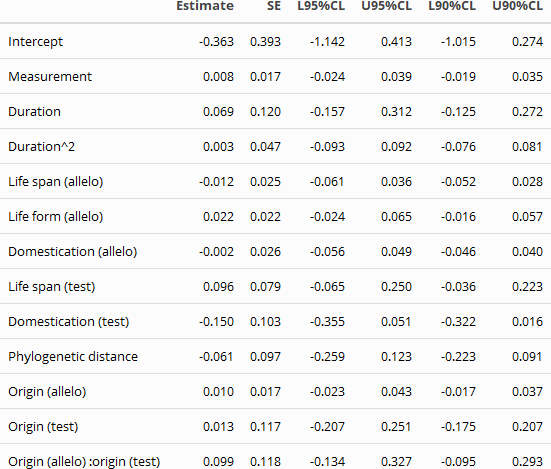

**Table S8** Effects of study design, biological traits and evolutionary history on effect sizes (ln RR) of allelopathy in **activated carbon** method.

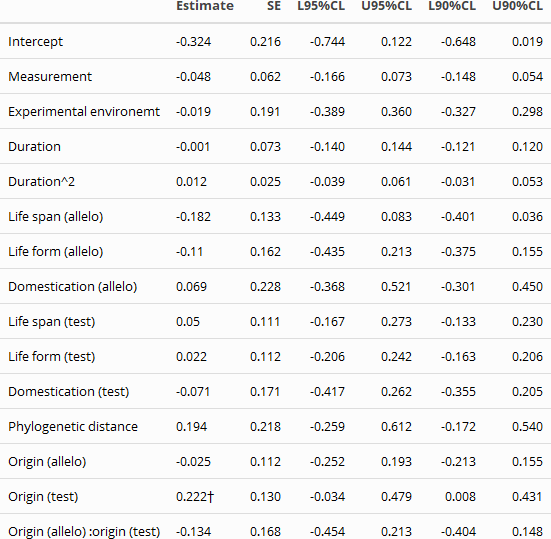

**Table S9** Effects of study design, biological traits and evolutionary history on effect sizes (ln RR) of allelopathy in **solvent extraction** method.

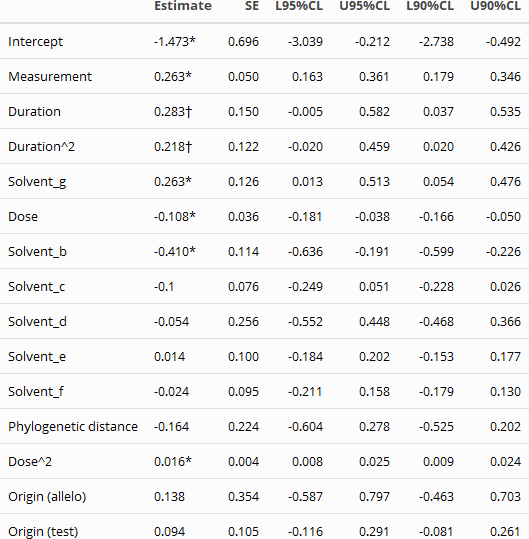

Types of solvents (from low to high polarity): a: Hexane (reference level); b: Diethyl ether; c: Ethyl acetate; d: Dichloromethane; e: n-Butanol; f: Methanol; g: water (the remaining fraction after extracted by organic solvents).

### Supplement S4 The complete funnel plot.

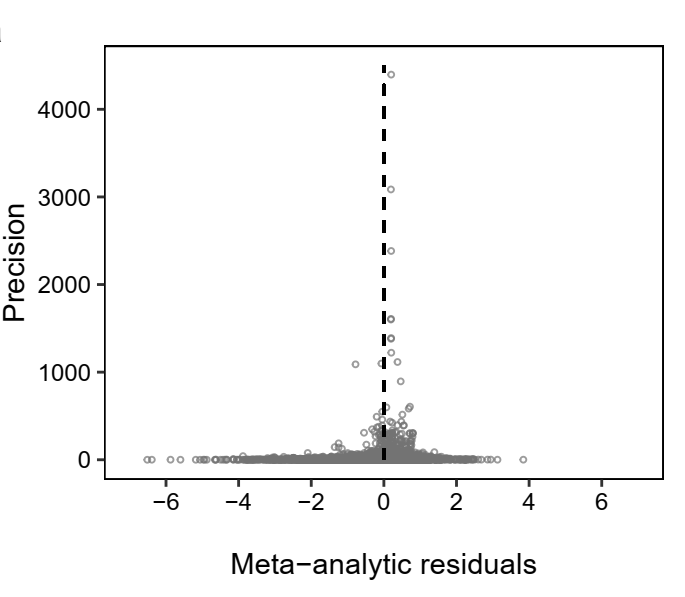

**Figure S3** Funnel plot of the study precision (inverse of standard error) *vs.* the residuals of the meta-analysis.

### Supplement S5 The phylogenetic tree for all species included in the meta-analysis

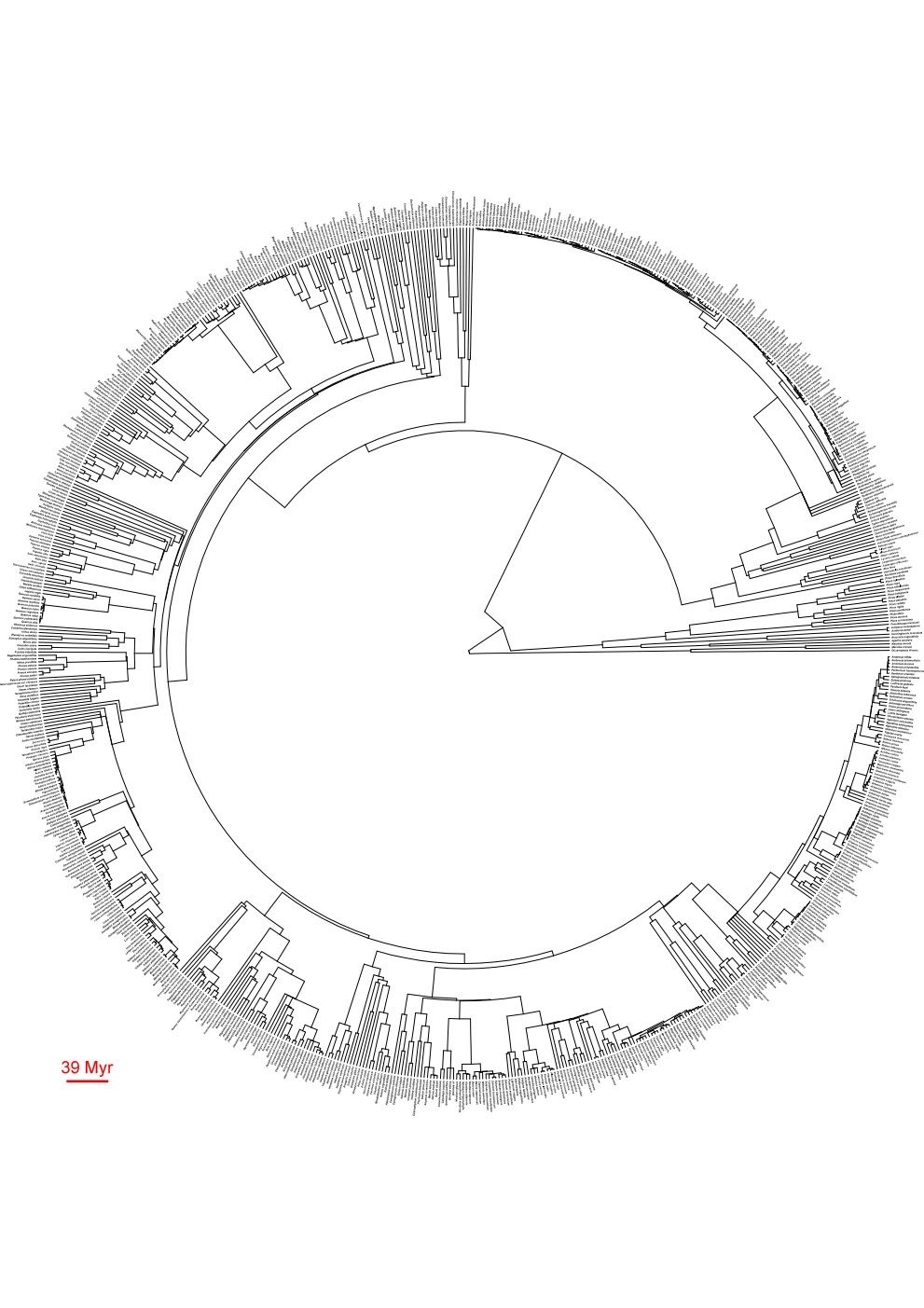

### Supplement S6 The 379 published studies that were included in the database

^1-226227-378^

1	Abdel-Farid, I., El-Sayed, M. & Mohamed, E. Allelopathic Potential of Calotropis procera and Morettia philaeana. *Int. J. Agric. Biol.* **15**, 130-134 (2013).

2	Aguilera, N. *et al.* Allelopathic effect of the invasive Acacia dealbata Link (Fabaceae) on two native plant species in south-central Chile. *Gayana Bot.* **72**, 231-239 (2015).

3	Aguilera, N. *et al.* Effects and identification of chemical compounds released from the invasive Acacia dealbata Link. *Chem. Ecol.* **31**, 479-493 (2015).

4	Al Harun, M. A., Johnson, J., Uddin, M. N. & Robinson, R. W. Identification and phytotoxicity assessment of phenolic compounds in Chrysanthemoides monilifera subsp. Monilifera (boneseed). *PLoS One* **10**, e0139992 (2015).

5	Ali, H. H. *et al.* Allelopathic effects of Rhynchosia capitata on germination and seedling growth of mungbean. *Planta Daninha* **31**, 501-509 (2013).

6	Al-Johani, N. S., Aytah, A. A. & Boutraa, T. Allelopathic impact of two weeds, Chenopodium murale and Malva parviflora on growth and photosynthesis of barley (Hordeum vulgare l.). *Pak. J. Bot.* **44**, 1865-1872 (2012).

7	Al-Sherif, E., Hegazy, A. K., Gomaa, N. H. & Hassan, M. Q. Allelopathic Effect of Black Mustard Tissues and Root Exudates on Some Crops and Weeds. *Planta Daninha* **31**, 11-19 (2013).

8	Álvarez-Iglesias, L., Puig, C. G., Garabatos, A., Reigosa, M. J. & Pedrol, N. Vicia faba aqueous extracts and plant material can suppress weeds and enhance crops. *Allelopathy J.* **34**, 299-314 (2014).

9	Amini, R., An, M., Pratley, J. & Azimi, S. Allelopathic assesment of annual ryegrass (Lolium rigidum): Bioassays. *Allelopathy J.* **24**, 67-76 (2009).

10	Amoo, S. O., Jo, A. U. & Van Staden, J. Allelopathic potential of Tetrapleura tetraptera leaf extracts on early seedling growth of five agricultural crops. *S. Afr. J. Bot.* **74**, 149-152 (2008).

11	Andrade, H. M. d., Bittencourt, A. H. C. & Vestena, S. Potencial alelopático de Cyperus rotundus L. sobre espécies cultivadas. *Ciência e Agrotecnologia* **33**, 1984-1990 (2009).

12	Araniti, F., Lupini, A., Sorgona, A., Statti, G. A. & Abenavoli, M. R. Phytotoxic activity of foliar volatiles and essential oils of Calamintha nepeta (L.) Savi. *Nat Prod Res* **27**, 1651-1656 (2013).

13	Araniti, F., Lupini, A., Sunseri, F. & Abenavoli, M. R. Allelopatic potential of Dittrichia viscosa (l.) w. Greuter mediated by vocs: A physiological and metabolomic approach. *PLoS One* **12**, e0170161 (2017).

14	Araniti, F., Sunseri, F. & Abenavoli, M. R. Phytotoxic activity and phytochemical characterization of Lotus ornithopodioides L., a spontaneous species of Mediterranean area. *Phytochem. Lett.* **8**, 179-183 (2014).

15	Arowosegbe, S. & Afolayan, A. J. Assessment of allelopathic properties of Aloe ferox Mill. on turnip, beetroot and carrot. *Biol Res* **45**, 363-368 (2012).

16	Aslam, F. *et al.* Differential Allelopathic Activity of Parthenium Hysterophorus L. Against Canary Grass and Wild Oat. *J. Anim. Plant Sci.* **24**, 234-244 (2014).

17	Aslam, F., Khaliq, A., Tanveer, A. & Zahir, Z. A. Wheat Herbage Amendments Alter Emergence Dynamics, Seedling Growth of Lambsquarter and Soil Properties. *Planta Daninha* **33**, 643-662 (2015).

18	Aslam, F., Khaliq, A., Tanveer, A., Zahir, Z. A. & Matloob, A. Wheat Residue Incorporation Modulate Emergence and Seedling Growth of Canary Grass by Affecting Biochemical Attributes and Soil Properties. *Int. J. Agric. Biol.* **18**, 1033-1042 (2016).

19	Aslani, F., Juraimi, A. S., Ahmad-Hamdani, M. S., Hashemi, F. S. G. & Alam, M. A. Control of weeds in glasshouse and rice field conditions by phytotoxic effects of Tinospora crispa (L.) Hook. f. & Thomson leaves. *Chil. J. Agric. Res.* **76**, 9 (2016).

20	Aslani, F. *et al.* Allelopathic effect of methanol extracts from Tinospora tuberculata on selected crops and rice weeds. *Acta Agric. Scand. Sect. B-Soil Plant Sci.* **64**, 165-177 (2014).

21	Aziz, S. & Shaukat, S. S. Degradation of phenolics in Digera muricata: Phytotoxic effects of root and shoot leachate plus N fertilization on the growth of millet. *Pak. J. Bot.* **47**, 287-290 (2015).

22	Bai, R., Zhao, X., Ma, F. & Li, C. Identification and bioassay of allelopathic substances from the root exudates of Mains prunifolia. *Allelopathy J.* **23**, 477-484 (2009).

23	Bainard, L. D., Brown, P. D. & Upadhyaya, M. K. Inhibitory effect of tall hedge mustard (Sisymbrium loeselii) allelochemicals on rangeland plants and arbuscular mycorrhizal fungi. *Weed Sci.* **57**, 386-393 (2009).

24	Bajwa, R. & Naz, I. Allelopathic effect of Eucalyptus citriodora on growth, nodulation and AM colonization of Vigna radiata (L) Wilczek. *Allelopathy J.* **15**, 237-246 (2005).

25	Baležentienė, L. Secondary metabolite accumulation and phytotoxicity of invasive species Solidago canadensis L. during the growth period. *Allelopathy J.* **35**, 217-226 (2015).

26	Balezentiene, L. & Renco, M. Phytotoxicity and accumulation of secondary metabolites in Heracleum mantegazzianum (Apiaceae). *Allelopathy J.* **33**, 267-276 (2014).

27	Bali, A. S., Batish, D. R., Singh, H. P., Kaur, S. & Kohli, R. K. Phytotoxicity and weed management potential of leaf extracts of Callistemon viminalis against the weeds of rice. *Acta Physiol. Plant.* **39**, 9 (2017).

28	Barritt, A. R. & Facelli, J. M. Effects of Casuarina pauper litter and grove soil on emergence and growth of understorey species in arid lands of South Australia. *J. Arid. Environ.* **49**, 569-579 (2001).

29	Barto, E. K. & Cipollini, D. Garlic Mustard (Alliaria petiolata) Removal Method Affects Native Establishment. *Invasive Plant Sci. Manag.* **2**, 230-236 (2009).

30	Barto, E. K. & Cipollini, D. Density-dependent phytotoxicity of impatiens pallida plants exposed to extracts of Alliaria petiolata. *J Chem Ecol* **35**, 495-504 (2009).

31	Barto, K., Friese, C. & Cipollini, D. Arbuscular mycorrhizal fungi protect a native plant from allelopathic effects of an invader. *J Chem Ecol* **36**, 351-360 (2010).

32	Basotra, R., Chauhan, S. & Todaria, N. P. Allelopathic effects of medicinal plants on food crops in Garhwal, Himalaya. *J. Sustain. Agric.* **26**, 43-56 (2005).

33	Batish, D. R., Kaur, M., Singh, H. P. & Kohli, R. K. Phytotoxicity of a medicinal plant, Anisomeles indica, against Phalaris minor and its potential use as natural herbicide in wheat fields. *Crop Prot.* **26**, 948-952 (2007).

34	Batish, D. R., Kaur, S., Singh, H. P. & Kohli, R. K. Nature of interference potential of leaf debris of Ageratum conyzoides. *Plant Growth Regul.* **57**, 137-144 (2009).

35	Batish, D. R., Kaur, S., Singh, H. P. & Kohli, R. K. Role of root-mediated interactions in phytotoxic interference of Ageratum conyzoides with rice (Oryza sativa). *Flora* **204**, 388-395 (2009).

36	Batish, D. R., Lavanya, K., Singh, H. P. & Kohli, R. K. Phenolic allelochemicals released by Chenopodium murale affect the growth, nodulation and macromolecule content in chickpea and pea. *Plant Growth Regul.* **51**, 119-128 (2007).

37	Batish, D. R., Lavanya, K., Singh, H. P. & Kohli, R. K. Root-mediated allelopathic interference of nettle-leaved goosefoot (Chenopodium murale) on wheat (Triticum aestivum). *Journal of Agronomy and Crop Science* **193**, 37-44 (2007).

38	Batish, D. R., Tung, P., Singh, H. P. & Kohli, R. K. Phytotoxicity of sunflower residues against some summer season crops. *Journal of Agronomy and Crop Science* **188**, 19-24 (2002).

39	Bauer, J. T., Shannon, S. M., Stoops, R. E. & Reynolds, H. L. Context dependency of the allelopathic effects of Lonicera maackii on seed germination. *Plant Ecol.* **213**, 1907-1916 (2012).

40	Bennett, A. E., Thomsen, M. & Strauss, S. Y. Multiple mechanisms enable invasive species to suppress native species. *Am J Bot* **98**, 1086-1094 (2011).

41	Bertin, C., Paul, R. N., Duke, S. O. & Weston, L. A. Laboratory assessment of the allelopathic effects of fine leaf fescues. *J Chem Ecol* **29**, 1919-1937 (2003).

42	Bhatt, B. P., Kumar, M. & Todaria, N. P. Studies on the allelopathic effects of terminalia species of Garhwal Himalaya. *J. Sustain. Agric.* **11**, 71-84 (1997).

43	Bilalis, D. J. *et al.* Evaluation of the allelopathic potential of quinoa (Chenopodium quinoa willd.). *Rom. Agric. Res.* **30**, 359-364 (2013).

44	Bogatek, R., Gniazdowska, A., Zakrzewska, W., Oracz, K. & Gawroński, S. W. Allelopathic effects of sunflower extracts on mustard seed germination and seedling growth. *Biologia Plantarum* **50**, 156-158 (2006).

45	Bossdorf, O., Shuja, Z. & Banta, J. A. Genotype and maternal environment affect belowground interactions between Arabidopsis thaliana and its competitors. *Oikos* **118**, 1541-1551 (2009).

46	Bousquet-Melou, A. *et al.* Allelopathic potential of Medicago arborea, a Mediterranean invasive shrub. *Chemoecology* **15**, 193-198 (2005).

47	Braine, J. W., Curcio, G. R., Wachowicz, C. M. & Hansel, F. A. Allelopathic effects of Araucaria angustifolia needle extracts in the growth of Lactuca sativa seeds. *J Forest Res-Jpn* **17**, 440-445 (2012).

48	Butnariu, M. An analysis of Sorghum halepense's behavior in presence of tropane alkaloids from Datura stramonium extracts. *Chem Cent J* **6**, 75 (2012).

49	Cai, S. L. & Mu, X. Q. Allelopathic potential of aqueous leaf extracts of Datura stramonium L. on seed germination, seedling growth and root anatomy of Glycine max (L.) Merrill. *Allelopathy J.* **30**, 235-246 (2012).

50	Callaway, R. M. & Aschehoug, E. T. Invasive plants versus their new and old neighbors: a mechanism for exotic invasion. *Science* **290**, 521-523 (2000).

51	Callaway, R. M., Ridenour, W. M., Laboski, T., Weir, T. & Vivanco, J. M. Natural selection for resistance to the allelopathic effects of invasive plants. *J. Ecol.* **93**, 576-583 (2005).

52	Cavieres, L. A., Chacon, P., Penaloza, A., Molina-Montenegro, M. A. & Arroyo, M. T. K. Leaf litter of Kageneckia angustifolia D. Don (Rosaceae) inhibits seed germination in sclerophyllous montane woodlands of central Chile. *Plant Ecol.* **190**, 13-22 (2007).

53	Chai, T. T., Ngoi, J. C. & Wong, F. C. Herbicidal Potential of Eichhornia crassipes Leaf Extract against Mimosa pigra and Vigna radiata. *Int. J. Agric. Biol.* **15**, 835-842 (2013).

54	Chapla, T. E. & Campos, J. B. Allelopathic Evidence in Exotic Guava (Psidium guajava L.). *Braz. Arch. Biol. Technol.* **53**, 1359-1362 (2010).

55	Chaves, N. & Escudero, J. C. Allelopathic effect of Cistus ladanifer on seed germination. *Funct. Ecol.* **11**, 432-440 (1997).

56	Chaves, N., Sosa, T., Alias, J. C. & Escudero, J. C. Germination inhibition of herbs in Cistus landanifer L. soils: Possible involvement of allelochemicals. *Allelopathy J.* **11**, 31-42 (2003).

57	Chen, F., Peng, S., Chen, B., Ni, G. Y. & Liao, H. X. Allelopathic potential and volatile compounds of Rosmarinus officinalis L. against weeds. *Allelopathy J.* **32**, 57-66 (2013).

58	Chen, H. *et al.* Growth and developmental responses of spinach (Spinacia oleracea L.) to the decomposing leaf litter of blue gum (Eucalyptus maidenii F. Muell.). *Allelopathy J.* **34**, 33-48 (2014).

59	Chen, S. Y., Xiao, S. & Callaway, R. M. Light intensity alters the allelopathic effects of an exotic invader. *Plant Ecol. Divers.* **5**, 521-526 (2012).

60	Chon, S. U. *et al.* Effects of alfalfa leaf extracts and phenolic allelochemicals on early seedling growth and root morphology of alfalfa and barnyard grass. *Crop Prot.* **21**, 1077-1082 (2002).

61	Chon, S. U. & Kim, J. D. Biological activity and quantification of suspected allelochemicals from alfalfa plant parts. *Journal of Agronomy and Crop Science* **188**, 281-285 (2002).

62	Christina, M. *et al.* Allelopathic effect of a native species on a major plant invader in Europe. *Naturwissenschaften* **102**, 12 (2015).

63	Chuah, T. S., Tiun, S. M. & Ismail, B. S. Allelopathic potential of crops on germination and growth of goosegrass (Eleusine indica L. Gaertn) weed. *Allelopathy J.* **27**, 33-42 (2011).

64	Ciarka, D., Gawronska, H., Malecka, M. & Gawronski, S. W. Allelopathic potential of sunflower. II. Allelopathic activity of plants compounds released in environment. *Allelopathy J.* **23**, 243-254 (2009).

65	Cipollini, D. & Darning, M. Direct and indirect effects of conditioned soils and tissue extracts of the invasive shrub, Lonicera maackii, on target plant performance. *Castanea* **73**, 166-176 (2008).

66	Cipollini, D., Stevenson, R. & Cipollini, K. Contrasting effects of allelochemicals from two invasive plants on the performance of a nonmycorrhizal plant. *Int. J. Plant Sci.* **169**, 371-375 (2008).

67	Cipollini, D., Stevenson, R., Enright, S., Eyles, A. & Bonello, P. Phenolic metabolites in leaves of the invasive shrub, Lonicera maackii, and their potential phytotoxic and anti-herbivore effects. *J Chem Ecol* **34**, 144-152 (2008).

68	Cipollini, K. A. & Schradin, K. D. Guilty in the court of public opinion: Testing presumptive impacts and allelopathic potential of Ranunculus ficaria. *American Midland Naturalist* **166**, 63-74 (2011).

69	Corbett, B. F. & Morrison, J. A. The allelopathic potentials of the non-native invasive plant Microstegium vimineum and the native Ageratina altissima: Two dominant species of the eastern forest herb layer. *Northeast. Nat* **19**, 297-312 (2012).

70	Correa, L. R., Soares, G. L. G. & Fett-Neto, A. G. Allelopathic potential of Psychotria leiocarpa, a dominant understorey species of subtropical forests. *S. Afr. J. Bot.* **74**, 583-590 (2008).

71	Cruz-Ortega, R., Álvarez-Añorve, M., Romero-Romero, M. T., Lara-Nunez, A. & Anaya, A. L. Growth and oxidative damage effects of Sicyos deppei weed on tomato. *Allelopathy J.* **21**, 83-94 (2008).

72	Cui, C., Cai, J., Jiang, Z. M. & Zhang, S. Effects of walnut (Juglans regia L.) root exudates on germination, seed growth and enzymatic activities of turnip (Brassica rapa L.). *Allelopathy J.* **28**, 237-250 (2011).

73	Cummings, J. A., Parker, I. M. & Gilbert, G. S. Allelopathy: a tool for weed management in forest restoration. *Plant Ecol.* **213**, 1975-1989 (2012).

74	Dadkhah, A. Phytotoxic potential of sugar beet (Beta vulgaris) and eucalyptus (Eucalyptus camaldulensis) to control purslane (Portulaca oleracea) weed. *Acta Agric. Scand. Sect. B-Soil Plant Sci.* **63**, 46-51 (2013).

75	Dai, Z. C. *et al.* Effects of leaf litter on inter-specific competitive ability of the invasive plant Wedelia trilobata. *Ecol. Res.* **31**, 367-374 (2016).

76	Dandelot, S., Robles, C., Pech, N., Cazaubon, A. & Verlaque, R. Allelopathic potential of two invasive alien Ludwigia spp. *Aquat. Bot.* **88**, 311-316 (2008).

77	Del Fabbro, C. & Prati, D. Invasive plant species do not create more negative soil conditions for other plants than natives. *Perspectives in Plant Ecology, Evolution and Systematics* **17**, 87-95 (2015).

78	Dheeba, B., Sampathkumar, P., Kannan, K., Kannan, M. & Tamizharasi, T. Comparison of herbicidal activity of Datura metal and Nerium oleander on the weed, Parthenium hysterophorus in green gram crop. *Natl. Acad. Sci. Lett.-India* **37**, 269-274 (2014).

79	Dias, A. S. & Dias, L. S. Interactions in allelopathic effects of Raphanus raphanistrum L. *Allelopathy J.* **19**, 495-500 (2007).

80	Dommanget, F. *et al.* Differential allelopathic effects of Japanese knotweed on willow and cottonwood cuttings used in riverbank restoration techniques. *J Environ Manage* **132**, 71-78 (2014).

81	Donnelly, M. J., Green, D. M. & Walters, L. J. Allelopathic effects of fruits of the Brazilian pepper Schinus terebinthifolius on growth, leaf production and biomass of seedlings of the red mangrove Rhizophora mangle and the black mangrove Avicennia germinans. *J. Exp. Mar. Biol. Ecol.* **357**, 149-156 (2008).

82	Dorning, M. & Cipollini, D. Leaf and root extracts of the invasive shrub, Lonicera maackii, inhibit seed germination of three herbs with no autotoxic effects. *Plant Ecol.* **184**, 287-296 (2006).

83	Dragoeva, A. P., Nanova, Z. D. & Kalcheva, V. P. Allelopathic activity of micropropagated Hyssopus officinalis L., Lamiaceae, water infusions. *Brazilian Journal of Pharmacognosy* **20**, 513-518 (2009).

84	Economou, G., Tzakou, O., Gani, A., Yannitsaros, A. & Bilalis, D. Allelopathic effect of Conyza albida on Avena sativa and Spirodela polyrhiza. *Journal of Agronomy and Crop Science* **188**, 248-253 (2002).

85	El Id, V. L. *et al.* Phytotoxic effect of Sesbania virgata (Cav.) Pers. on seeds of agronomic and forestry species. *J. For. Res.* **26**, 339-346 (2015).

86	El-Keblawy, A. & Abdelfatah, M. A. Impacts of native and invasive exotic Prosopis congeners on soil properties and associated flora in the arid United Arab Emirates. *J. Arid. Environ.* **100**, 1-8 (2014).

87	El-Kenany, E. T., El-Darier, S. M., Abdellatif, A. A. & Shaklol, S. M. Allelopathic potential of invasive species: Nicotiana glauca Graham on some ecological and physiological aspects of Medicago sativa L. and Triticum aestivum L. *Rend. Lincei.-Sci. Fis. Nat.* **28**, 159-167 (2017).

88	Endeshaw, S. T., Lodolini, E. M. & Neri, D. Effects of olive shoot residues on shoot and root growth of potted olive plantlets. *Sci. Hortic.* **182**, 31-40 (2015).

89	Ens, E. J., French, K., Bremner, J. B. & Korth, J. Novel technique shows different hydrophobic chemical signatures of exotic and indigenous plant soils with similar effects of extracts on indigenous species seedling growth. *Plant Soil* **326**, 403-414 (2010).

90	Eom, S. H., Yang, H. S. & Weston, L. A. An evaluation of the allelopathic potential of selected perennial groundcovers: foliar volatiles of catmint (Nepeta x faassenii) inhibit seedling growth. *J Chem Ecol* **32**, 1835-1848 (2006).

91	Eppard, H. R., Horton, J. L., Nilsen, E. T., Galusky, P. & Clinton, B. D. Investigating the allelopathic potential of Kalmia latifolia L. (Ericaceae). *Southeastern Naturalist* **4**, 383-392 (2005).

92	Erez, M. E. & Fİdan, M. Allelopathic effects of Sage (Salvia macrochlamys) extract on germination of Portulaca oleracea seeds. *Allelopathy J.* **35**, 285-296 (2015).

93	Escudero, A., Albert, M. J., Pita, J. M. & Perez-Garcia, F. Inhibitory effects of Artemisia herba-alba on the germination of the gypsophyte Helianthemum squamatum. *Plant Ecol.* **148**, 71-80 (2000).

94	Fan, L., Chen, Y., Yuan, J. G. & Yang, Z. Y. The effect of Lantana camara Linn. invasion on soil chemical and microbiological properties and plant biomass accumulation in southern China. *Geoderma* **154**, 370-378 (2010).

95	Fang, B. *et al.* Allelopathic effects of Eucalyptus urophylla on ten tree species in south China. *Agroforestry Systems* **76**, 401-408 (2008).

96	Farooq, M., Hussain, T., Wakeel, A. & Cheema, Z. A. Differential Response of Maize and Mungbean to Tobacco Allelopathy. *Exp. Agric.* **50**, 611-624 (2014).

97	Farooq, M., Jabran, K., Rehman, H. A. & Hussain, M. Allelopathic effects of rice on seedling development in wheat, oat, barley and berseem. *Allelopathy J.* **22**, 385-390 (2008).

98	Fernandez, C. *et al.* Potential allelopathic effect of Pinus halepensis in the secondary succession: an experimental approach. *Chemoecology* **16**, 97-105 (2006).

99	Fernandez, C. *et al.* The Impact of Competition and Allelopathy on the Trade-Off between Plant Defense and Growth in Two Contrasting Tree Species. *Front Plant Sci* **7**, 594 (2016).

100	Fernandez, C. *et al.* Regeneration failure of Pinus halepensis Mill.: The role of autotoxicity and some abiotic environmental parameters. *For. Ecol. Manage.* **255**, 2928-2936 (2008).

101	Filep, R. *et al.* Can seasonal dynamics of allelochemicals play a role in plant invasions? A case study with Helianthus tuberosus L. *Plant Ecol.* **217**, 1489-1501 (2016).

102	Foletto, M. P. *et al.* Allelopathic effects of Brachiaria ruziziensis and aconitic acid on Ipomoea triloba weed. *Allelopathy J.* **30**, 33-47 (2012).

103	Furness, N. H., Adomas, B., Dai, Q. J., Li, S. X. & Upadhyaya, M. K. Allelopathic influence of houndstongue (Cynoglossum officinale) and its modification by UV-B radiation. *Weed Technol.* **22**, 101-107 (2008).

104	Gallet, C. Allelopathic potential in bilberry-spruce forests: Influence of phenolic compounds on spruce seedlings. *J. Chem. Ecol.* **20**, 1009-1024 (1994).

105	Garnett, E., Jonsson, L. M., Dighton, J. & Murnen, K. Control of pitch pine seed germination and initial growth exerted by leaf litters and polyphenolic compounds. *Biol. Fertil. Soils* **40**, 421-426 (2004).

106	Gatti, A. B., Ferreira, A. G., Arduin, M. & Perez, S. C. G. d. A. Allelopathic effects of aqueous extracts of Artistolochia esperanzae O.Kuntze on development of Sesamum indicum L. seedlings. *Acta Bot. Bras.* **24**, 454-461 (2010).

107	Ghebrehiwot, H. M., Aremu, A. O. & Van Staden, J. Evaluation of the allelopathic potential of five South African mesic grassland species. *Plant Growth Regul.* **72**, 155-162 (2014).

108	Giordano, S. *et al.* In vitro allelopathic properties of wild rocket (Diplotaxis tenuifolia DC) extract and of its potential allelochemical S-glucopyranosyl thiohydroximate. *J. Plant Interact.* **1**, 51-60 (2005).

109	Gniazdowska, A., Oracz, K. & Bogatek, R. Phytotoxic effects of sunflower (Helianthus annuus L.) leaf extracts on germinating mustard (Sinapis alba L.) seeds. *Allelopathy J.* **19**, 215-226 (2007).

110	Golisz, A., Gawronska, H. & Gawronski, S. W. Influence of buckwheat allelochemicals on crops and weeds. *Allelopathy J.* **19**, 337-350 (2007).

111	Gomaa, N. H. & AbdElgawad, H. R. Phytotoxic effects of Echinochloa colona (L.) Link. (Poaceae) extracts on the germination and seedling growth of weeds. *Span. J. Agric. Res.* **10**, 492-501 (2012).

112	Goncalves, S., Ferraz, M. & Romano, A. Phytotoxic properties of Drosophyllum lusitanicum leaf extracts and its main compound plumbagin. *Sci. Hortic.* **122**, 96-101 (2009).

113	Goodall, J., Witkowski, E. T. F., Ammann, S. & Reinhardt, C. Does allelopathy explain the invasiveness of Campuloclinium macrocephalum (pompom weed) in the South African grassland biome? *Biol. Invasions* **12**, 3497-3512 (2010).

114	Grant, D. W., Peters, D. P. C., Beck, G. K. & Fraleigh, H. D. Influence of an exotic species, Acroptilon repens (L.) DC. on seedling emergence and growth of native grasses. *Plant Ecol.* **166**, 157-166 (2003).

115	Greer, M. J., Wilson, G. W. T., Hickman, K. R. & Wilson, S. M. Experimental evidence that invasive grasses use allelopathic biochemicals as a potential mechanism for invasion: chemical warfare in nature. *Plant Soil* **385**, 165-179 (2014).

116	Grisi, P. U., Gualtieri, S. C. J., Ranal, M. A. & Santana, D. G. Allelopathic interference of Sapindus saponaria root and mature leaf aqueous extracts on diaspore germination and seedling growth of Lactuca sativa and Allium cepa. *Braz. J. Bot.* **35**, 1-9 (2012).

117	Grove, S., Haubensak, K. A. & Parker, I. M. Direct and indirect effects of allelopathy in the soil legacy of an exotic plant invasion. *Plant Ecol.* **213**, 1869-1882 (2012).

118	Gruntman, M., Zieger, S. & Tielborger, K. Invasive success and the evolution of enhanced weaponry. *Oikos* **125**, 59-65 (2016).

119	Guerrero, P. C. & Bustamante, R. O. Can native tree species regenerate in Pinus radiata plantations in Chile? *For. Ecol. Manage.* **253**, 97-102 (2007).

120	Gulzar, A. & Siddiqui, M. B. Root-mediated allelopathic interference of bhringraj (Eclipta alba L.) Hassk. on peanut (Arachis hypogaea) and mung bean (Vigna radiata). *Appl. Soil Ecol.* **87**, 72-80 (2015).

121	Gunarathne, R. & Perera, G. A. D. Does the invasion of Prosopis juliflora cause the die-back of the native Manilkara hexandra in seasonally dry tropical forests of Sri Lanka? *Trop. Ecol.* **57**, 475-488 (2016).

122	Gupta, K., Jain, V., Solanki, I. S. & Tulika. Effect of aqueous extracts of root and stubble of oat (Avena sativa L.) on seedling growth and protein utilization in mung bean (Vigna radiata L.). *Allelopathy J.* **16**, 279-287 (2005).

123	Habermann, E., Pontes, F. C., Pereira, V. C., Imatomi, M. & Gualtieri, S. C. Phytotoxic potential of young leaves from Blepharocalyx salicifolius (Kunth) O. Berg (Myrtaceae). *Braz J Biol* **76**, 531-538 (2016).

124	Han, X., Cheng, Z. H., Meng, H. W., Yang, X. L. & Ahmad, I. Allelopathic effect of decomposed garlic (Allium sativum l.) stalk on lettuce (l. Sativa var. Crispa l.). *Pak. J. Bot.* **45**, 225-233 (2013).

125	Harun, M. A. Y. A., Johnson, J. & Robinson, R. W. The contribution of volatilization and exudation to the allelopathic phytotoxicity of invasive Chrysanthemoides monilifera subsp. monilifera (boneseed). *Biol. Invasions* **17**, 3609-3624 (2015).

126	Harun, M. A. Y. A., Robinson, R. W., Johnson, J. & Uddin, M. N. Allelopathic potential of Chrysanthemoides monilifera subsp. monilifera (boneseed): A novel weapon in the invasion processes. *S. Afr. J. Bot.* **93**, 157-166 (2014).

127	Hassan, M. O. *et al.* Influence of Sonchus oleraceus L. Residue on Soil Properties and Growth of Some Plants. *Philipp. Agric. Sci.* **97**, 368-376 (2014).

128	He, H., Song, Q. M., Wang, Y. F. & Yu, S. X. Phytotoxic effects of volatile organic compounds in soil water taken from a Eucalyptus urophylla plantation. *Plant Soil* **377**, 203-215 (2014).

129	Hedenec, P. *et al.* Allelopathic effect of new introduced biofuel crops on the soil biota: A comparative study. *Eur. J. Soil Biol.* **63**, 14-20 (2014).

130	Herranz, J. M., Ferrandis, P., Copete, M. A., Duro, E. M. & Zalacaín, A. Effect of allelopathic compounds produced by Cistus ladanifer on germination of 20 Mediterranean taxa. *Plant Ecol.* **184**, 259-272 (2005).

131	Horiuchi, J. *et al.* The floral volatile, methyl benzoate, from snapdragon (Antirrhinum majus) triggers phytotoxic effects in Arabidopsis thaliana. *Planta* **226**, 1-10 (2007).

132	Horman, C. S. & Anderson, V. J. Understory species response to Utah juniper litter. *Journal of Range Management* **56**, 68-71 (2003).

133	Hou, Y.-P., Peng, S.-L., Chen, B.-M. & Ni, G.-Y. Inhibition of an invasive plant (Mikania micrantha H.B.K.) by soils of three different forests in lower subtropical China. *Biol. Invasions* **13**, 381-391 (2010).

134	Hovstad, K. A. & Ohlson, M. Conspecific versus heterospecific litter effects on seedling establishment. *Plant Ecol.* **204**, 33-42 (2009).

135	Hu, G. & Zhang, Z. H. Aqueous tissue extracts of Conyza canadensis inhibit the germination and shoot growth of three native herbs with no autotoxic effects. *Planta Daninha* **31**, 805-811 (2013).

136	Hu, G. & Zhang, Z. H. Allelopathic effects of Chromolaena odorata on native and non-native invasive herbs. *J. Food Agric. Environ.* **11**, 878-882 (2013).

137	Hu, G., Zhang, Z. H. & Hu, B. Q. in *Advances in Environmental Technologies, Pts 1-6* Vol. 726-731 *Advanced Materials Research* (eds J. Zhao, R. Iranpour, X. Li, & B. Jin) 4348-4351 (Trans Tech Publications Ltd, 2013).

138	Huang, W. *et al.* Impact of decomposing Cinnamomum septentrionale leaf litter on the growth of Eucalyptus grandis saplings. *Plant Physiol Biochem* **70**, 411-417 (2013).

139	Huang, W. W. *et al.* Impact of aqueous extracts of Cinnamomum septentrionale leaf litter on the growth and photosynthetic characteristics of Eucalyptus grandis seedlings. *New For.* **46**, 561-576 (2015).

140	Huang, Z., Haig, T., Wang, S. L. & Han, S. J. Autotoxicity of Chinese fir on seed germination and seedling growth. *Allelopathy J.* **9**, 187-193 (2002).

141	Hunter, M. E. & Menges, E. S. Allelopathic effects and root distribution of Ceratiola ericoides (Empetraceae) on seven rosemary scrub species. *Am J Bot* **89**, 1113-1118 (2002).

142	Hussain, M. I., Gonzalez, L. & Reigosa, M. J. Allelopathic potential of Acacia melanoxylon on the germination and root growth of native species. *Weed Biol. Manag.* **11**, 18-28 (2011).

143	Hussain, M. I., Gonzalez, L., Souto, C. & Reigosa, M. J. Ecophysiological responses of three native herbs to phytotoxic potential of invasive Acacia melanoxylon R. Br. *Agroforestry Systems* **83**, 149-166 (2011).

144	Imatomi, M., Novaes, P. & Gualtieri, S. C. J. Interspecific variation in the allelopathic potential of the family Myrtaceae. *Acta Bot. Bras.* **27**, 54-61 (2013).

145	Imatomi, M., Novaes, P., Miranda, M. A. F. M. & Gualtieri, S. C. J. Phytotoxic effects of aqueous leaf extracts of four Myrtaceae species on three weeds. *Acta Sci.-Agron.* **37**, 241-248 (2015).

146	Inderjit, Asakawa, C. & Dakshini, K. M. M. Allelopathic potential of Verbesina encelioides root leachate in soil. *Canadian Journal of Botany* **77**, 1419-1424 (2000).

147	Inderjit *et al.* Volatile chemicals from leaf litter are associated with invasiveness of a neotropical weed in Asia. *Ecology* **92**, 316-324 (2011).

148	Inderjit & Foy, C. L. Nature of the interference mechanism of mugwort (Artemisia vulgaris). *Weed Technol.* **13**, 176-182 (1999).

149	Iqbal, Z., Nasir, H., Hiradate, S. & Fujii, Y. Plant growth inhibitory activity of Lycoris radiata Herb. and the possible involvement of lycorine as an allelochemical. *Weed Biol. Manag.* **6**, 221-227 (2006).

150	Islam, A. K. & Kato-Noguchi, H. Phytotoxic activity of Ocimum tenuiflorum extracts on germination and seedling growth of different plant species. *ScientificWorldJournal* **2014**, 676242 (2014).

151	Islam, M. S., Iwasaki, A., Suenaga, K. & Kato-Noguchi, H. Isolation and identification of two potential phytotoxic substances from the aquatic fern Marsilea crenata. *J. Plant Biol.* **60**, 75-81 (2017).

152	Ismail, B. S., Tan, P. W. & Chuah, T. S. Assessment of the Potential Allelopathic Effects of Pennisetum purpureum Schumach. on the Germination and Growth of Eleusine indica (L.) Gaertn. *Sains Malays.* **44**, 269-274 (2015).

153	Jabeen, N., Ahmed, M., Shaukat, S. S. & Iram-us-Slam. Allelopathic Effects of Weeds on Wheat (Triticum Aestivum L.) Germination and Growth. *Pak. J. Bot.* **45**, 807-811 (2013).

154	Jandova, K., Dostal, P. & Cajthaml, T. Searching for Heracleum mantegazzianum allelopathy in vitro and in a garden experiment. *Biol. Invasions* **17**, 987-1003 (2015).

155	Jarchow, M. E. & Cook, B. J. Allelopathy as a mechanism for the invasion of Typha angustifolia. *Plant Ecol.* **204**, 113-124 (2009).

156	Javaid, A. & Riaz, T. Effects of application of green leaf manure on growth and mycorrhizal colonization of maize. *Allelopathy J.* **21**, 339-348 (2008).

157	Javaid, A., Shafique, S., Bajwa, R. & Shafique, S. Parthenium Management through Aqueous Extracts of Alstonia Scholaris. *Pak. J. Bot.* **42**, 3651-3657 (2010).

158	Javaid, A., Shafique, S. & Shafique, S. Herbicidal effects of extracts and residue incorporation of Datura metel against parthenium weed. *Nat Prod Res* **24**, 1426-1437 (2010).

159	Javaid, A., Shafique, S., Shafique, S. & Riaz, T. Effect of rice extracts and residue incorporation on Parthenium hysterophorus management. *Allelopathy J.* **22**, 353-262 (2008).

160	Jose, C. M. *et al.* Phytotoxic effects of phenolic acids from Merostachys riedeliana, a native and overabundant Brazilian bamboo. *Chemoecology* **26**, 235-246 (2016).

161	Kaligaric, M. *et al.* Grassland succession is mediated by umbelliferous colonizers showing allelopathic potential. *Plant Biosyst.* **145**, 688-698 (2011).

162	Kara, Y. & Kuru, A. Allelopathic effects of jojoba (Simmondsia chinensis) on seed germination and seedling growth of bean (Phaseolus vulgaris) and wheat (Triticum aestivum). *J. Environ. Prot. Ecol.* **16**, 588-593 (2015).

163	Kato-Noguchi, H. Effects of lemon balm (Melissa Officinalis L.) extract on germination and seedling growth of six plants. *Acta Physiol. Plant.* **23**, 49-53 (2001).

164	Kato-Noguchi, H. Assessment of the allelopathic potential of Ageratum conyzoides. *Biologia Plantarum* **44**, 309-311 (2001).

165	Kaya, M. D. *et al.* Allelopathic Role of Essential Oils in Sunflower Stubble on Germination and Seedling Growth of the Subsequent Crop. *Int. J. Agric. Biol.* **15**, 337-341 (2013).

166	Khaliq, A., Hussain, S., Matloob, A., Tanveer, A. & Aslam, F. Swine cress (Cronopus didymus l. Sm.) residues inhibit rice emergence and early seedling growth. *Philipp. Agric. Sci.* **96**, 419-425 (2013).

167	Khaliq, A., Hussain, S., Matloob, A., Wahid, A. & Aslam, F. Aqeous Swine Cress (Coronopus didymus) Extracts Inhibit Wheat Germination and Early Seedling Growth. *Int. J. Agric. Biol.* **15**, 743-748 (2013).

168	Khaliq, A., Matloob, A., Khan, M. B. & Tanveer, A. Differential Suppression of Rice Weeds by Allelopathic Plant Aqueous Extracts. *Planta Daninha* **31**, 21-28 (2013).

169	Khan, R. A. *et al.* Phytotoxic characterization of various fractions of Launaea procumbens. *African Journal of Biotechnology* **10**, 5377-5380 (2011).

170	Kim, Y. O., Johnson, J. D. & Lee, E. J. Phytotoxic effects and chemical analysis of leaf extracts from three Phytolaccaceae species in South Korea. *J Chem Ecol* **31**, 1175-1186 (2005).

171	Kim, Y. O. & Lee, E. J. Comparison of phenolic compounds and the effects of invasive and native species in East Asia: support for the novel weapons hypothesis. *Ecol. Res.* **26**, 87-94 (2011).

172	Kimura, F. & Kato-Noguchi, H. Allelopathic potential of Japanese red pine (Pinus densiflora Sieb. et Zucc.) needles and its litter. *Allelopathy J.* **32**, 123-132 (2013).

173	Ladwig, L. M., Meiners, S. J., Pisula, N. L. & Lang, K. A. Conditional allelopathic potential of temperate lianas. *Plant Ecol.* **213**, 1927-1935 (2012).

174	Lankau, R. Soil microbial communities alter allelopathic competition between Alliaria petiolata and a native species. *Biol. Invasions* **12**, 2059-2068 (2009).

175	Lara-Nunez, A., Sanchez-Nieto, S., Luisa Anaya, A. & Cruz-Ortega, R. Phytotoxic effects of Sicyos deppei (Cucurbitaceae) in germinating tomato seeds. *Physiol Plant* **136**, 180-192 (2009).

176	Laterra, P. & Bazzalo, M. E. Seed-to-seed allelopathic effects between two invaders of burned Pampa grasslands. *Weed Res.* **39**, 297-308 (1999).

177	Lau, J. A. *et al.* Inference of allelopathy is complicated by effects of activated carbon on plant growth. *New Phytol* **178**, 412-423 (2008).

178	Lawley, Y. E., Teasdale, J. R. & Weil, R. R. The Mechanism for Weed Suppression by a Forage Radish Cover Crop. *Agron. J.* **104**, 205-214 (2012).

179	Lehman, M. E. & Blum, U. Cover crop debris effects on weed emergence as modified by environmental factors. *Allelopathy J.* **4**, 69-88 (1997).

180	Lesica, P. & DeLuca, T. H. Is tamarisk allelopathic? *Plant Soil* **267**, 357-365 (2004).

181	Levizou, E., Karageorgou, P., Psaras, G. K. & Manetas, Y. Inhibitory effects of water soluble leaf leachates from Dittrichia viscosa on lettuce root growth, statocyte development and graviperception. *Flora* **197**, 152-157 (2002).

182	Li, G. *et al.* Allelopathic effects of decaying Italian ryegrass (Lolium mutiflorum Lam.) residues on rice. *Allelopathy J.* **22**, 15-24 (2008).

183	Li, J. M. & Jin, Z. X. Potential allelopathic effects of Mikania micrantha on the seed germination and seedling growth of Coix lacryma-jobi. *Weed Biol. Manag.* **10**, 194-201 (2010).

184	Li, X. F., Wang, J., Huang, D., Wang, L. X. & Wang, K. Allelopathic potential of Artemisia frigida and successional changes of plant communities in the northern China steppe. *Plant Soil* **341**, 383-398 (2010).

185	Li, X. X., Yu, M. F., Ruan, X., Zhang, Y. Z. & Wang, Q. Phytotoxicity of 4,8-dihydroxy-1-tetralone isolated from Carya cathayensis Sarg. to various plant species. *Molecules* **19**, 15452-15467 (2014).

186	Li, Y., Hu, T., Zeng, F., Chen, H. & Wu, X. Allelopathic effects of decomposing Eucalyptus grandis leaf litter on growth and physiology of Setaria viridis. *Allelopathy J.* **34**, 17-32 (2014).

187	Li, Y. B. & Zhang, Q. Effects of naturally and microbially decomposed cotton stalks on cotton seedling growth. *Arch. Agron. Soil Sci.* **62**, 1264-1270 (2016).

188	Li, Z. H. *et al.* Biologica activity and quantification of potential autotoxins from Picea schrenkiana leaves. *Allelopathy J.* **27**, 245-262 (2011).

189	Liu, L. Y., He, H. Z., Luo, S. M. & Li, H. S. Allelopathic potential of Rhus chinensis on seedling growth of radish, semen cassiae and black soyabean. *J. For. Res.* **26**, 273-279 (2015).

190	Liu, P. *et al.* Autotoxic potential of root exudates of peanut (Arachis hypogaea L.). *Allelopathy J.* **26**, 197-206 (2010).

191	Lodhi, M. A. K. & Rice, E. L. Allelopathic Effects of Celtis laevigata. *Bulletin of the Torrey Botanical Club* **98**, 83-89 (1971).

192	Lorenzo, P., Pazos-Malvido, E., Gonzalez, L. & Roger, M. J. Allelopathic interference of invasive Acacia dealbata: Physiological effects. *Allelopathy J.* **22**, 452-462 (2008).

193	Lorenzo, P., Pazos-Malvido, E., Reigosa, M. J. & González, L. Differential responses to allelopathic compounds released by the invasive Acacia dealbata Link (Mimosaceae) indicate stimulation of its own seed. *Aust. J. Bot.* **58**, 546–553 (2010).

194	Lorenzo, P. & Rodriguez-Echeverria, S. Influence of soil microorganisms, allelopathy and soil origin on the establishment of the invasive Acacia dealbata. *Plant Ecol. Divers.* **5**, 67-73 (2012).

195	Lou, Y., Davis, A. S. & Yannarell, A. C. Interactions between allelochemicals and the microbial community affect weed suppression following cover crop residue incorporation into soil. *Plant Soil* **399**, 357-371 (2016).

196	Loydi, A., Donath, T. W., Eckstein, R. L. & Otte, A. Non-native species litter reduces germination and growth of resident forbs and grasses: allelopathic, osmotic or mechanical effects? *Biol. Invasions* **17**, 581-595 (2015).

197	Ma, L. *et al.* Phytotoxic effects of Stellera chamaejasme L. root extract. *African Journal of Agricultural Research* **6**, 1170-1176 (2011).

198	Mahall, B. E. & Callaway, R. M. Root Communication Mechanisms and Intracommunity Distributions of 2 Mojave Desert Shrubs. *Ecology* **73**, 2145-2151 (1992).

199	Mahmood, K. *et al.* Exogenous application of signaling compounds enhances rice allelopathic potential in rhizosphere soil. *Int. J. Agric. Biol.* **15**, 1319-1324 (2013).

200	Mahmoud, A., Singh, S. D. & Muralikrishna, K. S. Allelopathy in jatropha plantation: Effects on seed germination, growth and yield of wheat in north-west India. *Agric. Ecosyst. Environ.* **231**, 240-245 (2016).

201	Mahmoud, T. S. *et al.* Biological activity of Gomphrena elegans Mart. Var. Elegans (Amaranthaceae). *Allelopathy J.* **28**, 201-212 (2011).

202	Mallik, A. U., Biswas, S. R. & Collier, L. C. S. Belowground interactions between Kalmia angustifolia and Picea mariana: roles of competition, root exudates and ectomycorrhizal association. *Plant Soil* **403**, 471-483 (2016).

203	Mallik, A. U. & Pellissier, F. Effects of Vaccinium myrtillus on spruce regeneration: Testing the notion of coevolutionary significance of allelopathy. *J. Chem. Ecol.* **26**, 2197-2209 (2000).

204	Mao, J. A. *et al.* Crude extract of Astragalus mongholicus root inhibits crop seed germination and soil nitrifying activity. *Soil Biol. Biochem.* **38**, 201-208 (2006).

205	McEwan, R. W., Arthur-Paratley, L. G., Rieske, L. K. & Arthur, M. A. A multi-assay comparison of seed germination inhibition by Lonicera maackii and co-occurring native shrubs. *Flora* **205**, 475-483 (2010).

206	Mecina, G. F. *et al.* Phytotoxicity of Tridax procumbens L. *S. Afr. J. Bot.* **102**, 130-136 (2016).

207	Mecina, G. F. *et al.* Phytotoxicity of extracts and fractions of Ouratea spectabilis (Mart. ex Engl.) Engl. (Ochnaceae). *S. Afr. J. Bot.* **95**, 174-180 (2014).

208	Mehmood, K., Asif, H. M., Bajwa, R., Shafique, S. & Shafique, S. Phytotoxic potential of bark extracts of Acacia nilotica and Syzygium cumini againstParthenium hysterophorus. *Pak. J. Bot.* **43**, 3007-3012 (2011).

209	Meksawat, S. & Pornprom, T. Allelopathic effect of itchgrass (Rottboellia cochinchinensis) on seed germination and plant growth. *Weed Biol. Manag.* **10**, 16-24 (2010).

210	Mengardo, A. L. T. & Pivello, V. R. The effects of an exotic palm on a native palm during the first demographic stages: contributions to ecological management. *Acta Bot. Bras.* **28**, 552-558 (2014).

211	Metlen, K. L., Aschehoug, E. T. & Callaway, R. M. Competitive outcomes between two exotic invaders are modified by direct and indirect effects of a native conifer. *Oikos* **122**, 632-640 (2013).

212	Metlen, K. L. & Callaway, R. M. Native North American pine attenuates the competitive effects of a European invader on native grasses. *Biol. Invasions* **17**, 1227-1237 (2015).

213	Mitic, N. *et al.* Use of Chenopodium murale L. transgenic hairy root in vitro culture system as a new tool for allelopathic assays. *J Plant Physiol* **169**, 1203-1211 (2012).

214	Mohamed, A. & El-Gawad, A. Ecology and allelopathic control of Brassica tournefortii in reclaimed areas of the Nile Delta, Egypt. *Turk. J. Bot.* **38**, 347-357 (2014).

215	Morgan, E. C. & Overholt, W. A. Potential allelopathic effects of Brazilian pepper (Schinus terebinthifolius Raddi, Anacardiaceae) aqueous extract on germination and growth of selected Florida native plants1. *The Journal of the Torrey Botanical Society* **132**, 11-15 (2005).

216	Mudrak, O. & Frouz, J. Allelopathic effect of Salix caprea litter on late successional plants at different substrates of post-mining sites: pot experiment studies. *Botany-Botanique* **90**, 311-318 (2012).

217	Navarro-Cano, J. A. Effect of grass litter on seedling recruitment of the critically endangered Cistus heterophyllus in Spain. *Flora* **203**, 663-668 (2008).

218	Navarro-Cano, J. A., Barbera, G. G. & Castillo, V. M. Pine Litter from Afforestations Hinders the Establishment of Endemic Plants in Semiarid Scrubby Habitats of Natura 2000 Network. *Restoration Ecology* **18**, 165-169 (2010).

219	Nazim, K., Ahmed, M., Shaukat, S. S., Khan, M. U. & Hussian, S. S. Auto toxicity of Avicennia marina (forsk.) vierh in pakistan. *Pak. J. Bot.* **46**, 465-470 (2014).

220	Nesic, M. *et al.* Allelopathic potential of the invasive species Aster lanceolatus Willd. *Period. Biol.* **118**, 1-7 (2016).

221	Ngondya, I. B., Munishi, L. K., Treydte, A. C. & Ndakidemi, P. A. A nature-based approach for managing the invasive weed species Gutenbergia cordifolia for sustainable rangeland management. *SpringerPlus* **5**, 1787 (2016).

222	Nickerson, K. & Flory, S. L. Competitive and allelopathic effects of the invasive shrub Schinus terebinthifolius (Brazilian peppertree). *Biol. Invasions* **17**, 555-564 (2015).

223	Nielsen, J. A., Frew, R. D., Whigam, P. A., Callaway, R. M. & Dickinson, K. J. M. Germination and growth responses of co-occurring grass species to soil from under invasive Thymus vulgaris. *Allelopathy J.* **35**, 139-152 (2015).

224	Nilsson, M. C. Separation of allelopathy and resource competition by the boreal dwarf shrub Empetrum hermaphroditum Hagerup. *Oecologia* **98**, 1-7 (1994).

225	Nilsson, M. C., Zackrisson, O., Sterner, O. & Wallstedt, A. Characterisation of the differential interference effects of two boreal dwarf shrub species. *Oecologia* **123**, 122-128 (2000).

226	Novoa, A., Gonzalez, L., Moravcova, L. & Pysek, P. Effects of soil characteristics, allelopathy and frugivory on establishment of the invasive plant Carpobrotus edulis and a co-occurring native, Malcolmia littorea. *PLoS One* **7**, e53166 (2012).

227	Oliveira, A. P. P., Pereira, S. R., CÂNdido, A. C. S., Laura, V. A. & Peres, M. T. L. P. Can Allelopathic Grasses Limit Seed Germination and Seedling Growth of Mutambo? A Test with Two Species of Brachiaria Grasses. *Planta Daninha* **34**, 639-648 (2016).

228	Oliveira, S. C. C. & Campos, M. L. Allelopathic effects of Solanum palinacanthum leaves on germination and seedling growth of Sesamum indicum. *Allelopathy J.* **18**, 331-338 (2006).

229	Oracz, K. *et al.* Induction of oxidative stress by sunflower phytotoxins in germinating mustard seeds. *J Chem Ecol* **33**, 251-264 (2007).

230	Orr, S. P., Rudgers, J. A. & Clay, K. Invasive plants can inhibit native tree seedlings: testing potential allelopathic mechanisms. *Plant Ecol.* **181**, 153-165 (2005).

231	Oveisi, M., Mashhadi, H. R., Baghestani, M. A., Alizadeh, H. M. & Badri, S. Assessment of the allelopathic potential of 17 Iranian barley cultivars in different development stages and their variations over 60 years of selection. *Weed Biol. Manag.* **8**, 225-232 (2008).

232	Overholt, W. A., Cuda, J. P. & Markle, L. Can novel weapons favor native plants? Allelopathic interactions betweenMorella cerifera(L.) andSchinus terebinthifoliaRaddi1. *The Journal of the Torrey Botanical Society* **139**, 356-366 (2012).

233	Ozpinar, H., Dag, S. & Yigit, E. Alleophatic effects of benzoic acid, salicylic acid and leaf extract of Persica vulgaris Mill. (Rosaceae). *S. Afr. J. Bot.* **108**, 102-109 (2017).

234	Padhy, B., Patnaik, P. K. & Tripathy, A. K. Allelopathic potential of Eucalyptus leaf litter leachates on germination and seedling growth of fingermillet. *Allelopathy J.* **7**, 69-78 (2000).

235	Parvez, S. S., Parvez, M. M. & Fujii, Y. Tamarindus indica L. leaf is a source of allelopathic substance. *Plant Growth Regul.* **40**, 107–115 (2003).

236	Pawlowski, A., Kaltchuk-Santos, E., Zini, C. A., Caramao, E. B. & Soares, G. L. G. Essential oils of Schinus terebinthifolius and S-molle (Anacardiaceae): Mitodepressive and aneugenic inducers in onion and lettuce root meristems. *S. Afr. J. Bot.* **80**, 96-103 (2012).

237	Peguero, G., Lanuza, O. R., Savé, R. & Espelta, J. M. Allelopathic potential of the neotropical dry-forest tree Acacia pennatula Benth.: inhibition of seedling establishment exceeds facilitation under tree canopies. *Plant Ecol.* **213**, 1945-1953 (2011).

238	Pilipavicius, V. Allelopathic effect of Elytrigia repens (L.) Nevski on germination and early growth of spring wheat. *J. Food Agric. Environ.* **10**, 1520-1523 (2012).

239	Pilipavicius, V., Romaneckas, K. & Cesna, J. Allelopathic effect of Sonchus arvensis L. on germination and early growth of spring wheat. *J. Food Agric. Environ.* **11**, 1060-1063 (2013).

240	Pisula, N. L. & Meiners, S. J. Relative allelopathic potential of invasive plant species in a young disturbed woodland1. *The Journal of the Torrey Botanical Society* **137**, 81-87 (2010).

241	Pollock, J. L., Seastedt, T. R., Callaway, R. M., Kaur, J. & Inderjit. Allelopathy and plant invasions: traditional, congeneric, and bio-geographical approaches. *Biol. Invasions* **10**, 875-890 (2008).

242	Prasad, S., Singh, B. & Todaria, N. P. Effects of Cassia tora on the germination and growth of Parthenium hysterophorus L. *Allelopathy J.* **17**, 303-310 (2006).

243	Prati, D. & Bossdorf, O. Allelopathic inhibition of germination by Alliaria petiolata (Brassicaceae). *Am J Bot* **91**, 285-288 (2004).

244	Preston, C. A., Betts, H. & Baldwin, I. T. Methyl jasmonate as an allelopathic agent: Sagebrush inhibits germination of a neighboring tobacco, Nicotiana Attenuata. *J. Chem. Ecol.* **28**, 2343-2369 (2002).

245	Prichoa, F. C., Leyser, G., de Oliveira, J. V. & Cansian, R. L. Comparative allelopathic effects of Cryptocarya moschata and Ocotea odorifera aqueous extracts on Lactuca sativa. *Acta Sci.-Agron.* **35**, 197-202 (2013).

246	Pudelko, K., Majchrzak, L. & Narozna, D. Allelopathic effect of fibre hemp (Cannabis sativa L.) on monocot and dicot plant species. *Ind. Crop. Prod.* **56**, 191-199 (2014).

247	Puig, C. G., Alvarez-Iglesias, L., Reigosa, M. J. & Pedrol, N. Eucalyptus globulus Leaves Incorporated as Green Manure for Weed Control in Maize. *Weed Sci.* **61**, 154-161 (2013).

248	Pula, J., Barabasz-Krasny, B., Mozdzen, K., Soltys-Lelek, A. & Lepiarczyk, A. Effect of aqueous extracts of sticky willy (Galium aparine l.) on the growth of seedlings of selected maize varieties (Zea mays l.). *Not. Bot. Horti Agrobot. Cluj-Na.* **44**, 518-524 (2016).

249	Qasem, J. R. Differences in the allelopathy results from field observations to laboratory and glasshouse experiment. *Allelopathy J.* **26**, 45-58 (2010).

250	Qin, B. *et al.* No evidence for root-mediated allelopathy in Centaurea solstitialis, a species in a commonly allelopathic genus. *Biol. Invasions* **9**, 897-907 (2007).

251	Quayyum, H. A., Mallik, A. U., Orr, D. E. & Le, P. F. Allelopathic potential of aquatic plants associated with wild rice: Ii. Isolation and identification of allelochemicals. *J. Chem. Ecol.* **25**, 221-228 (1999).

252	Quddus, M. S., Bellairs, S. M. & Wurm, P. A. S. Acacia holosericea (Fabaceae) litter has allelopathic and physical effects on mission grass (Cenchrus pedicellatus and C-polystachios) (Poaceae) seedling establishment. *Aust. J. Bot.* **62**, 189-195 (2014).

253	Rahman, A. Allelopathic potential of Parthenium hysterophorus L. on Cassia spp. *Allelopathy J.* **18**, 345-354 (2006).

254	Rashid, A., Furness, N. H., Ellis, B. E. & Upadhyaya, M. K. Inhibition of seed germination and seedling growth by hound’s-tongue (Cynoglossum officinale L.) seed leachate. *Weed Biol. Manag.* **5**, 143–149 (2005).

255	Rashid, I. & Reshi, Z. Allelopathic interaction of an alien invasive species Anthemis cotula on its neghbours Conyza canadensis and Galinosoga parviflora. *Allelopathy J.* **28**, 77-92 (2011).

256	Rashid, M. H., Asaeda, T. & Uddin, M. N. Litter-mediated allelopathic effects of kudzu (Pueraria montana) onBidens pilosaandLolium perenneand its persistence in soil. *Weed Biol. Manag.* **10**, 48-56 (2010).

257	Rashid, M. H., Asaeda, T. & Uddin, M. N. The Allelopathic Potential of Kudzu (Pueraria montana). *Weed Sci.* **58**, 47-55 (2010).

258	Rasmussen, J. A. & Rice, E. L. Allelopathic effects of Sporobolus pyramidatus on vegetational patterning. *American Midland Naturalist* **86**, 309-326 (1971).

259	Rawat, L. S., Narwal, S. S., Kadian, H. S. & Negi, V. S. Allelopathic effects of sunflower (Helianthus annuus) on germination and growth of Parthenium hysterophorus. *Allelopathy J.* **27**, 225-236 (2011).

260	Renne, I. J., Rios, B. G., Fehmi, J. S. & Tracy, B. F. Low allelopathic potential of an invasive forage grass on native grassland plants: a cause for encouragement? *Basic Appl. Ecol.* **5**, 261-269 (2004).

261	Rice, C. P., Park, Y. B., Adam, F., Abdul-Baki, A. A. & Teasdale, J. R. Hydroxamic acid content and toxicity of rye at selected growth stages. *J Chem Ecol* **31**, 1887-1905 (2005).

262	Rice, E. L. Allelopathic effects of Andropogon virginicus and its persistence in old fields. *Am. J. Bot.* **59**, 752-755 (1972).

263	Ridenour, W. M. & Callaway, R. M. The relative importance of allelopathy in interference: the effects of an invasive weed on a native bunchgrass. *Oecologia* **126**, 444-450 (2001).

264	Rivera-Vega, L. J. *et al.* Allelopathic effects of glucosinolate breakdown products in Hanza [Boscia senegalensis (Pers.) Lam.] processing waste water. *Front Plant Sci* **6**, 532 (2015).

265	Ruan, X. *et al.* Effects of climate warming on plant autotoxicity in forest evolution: a case simulation analysis for Picea schrenkiana regeneration. *Ecol Evol* **6**, 5854-5866 (2016).

266	Rudrappa, T., Bonsall, J., Gallagher, J. L., Seliskar, D. M. & Bais, H. P. Root-secreted allelochemical in the noxious weed Phragmites australis deploys a reactive oxygen species response and microtubule assembly disruption to execute rhizotoxicity. *J Chem Ecol* **33**, 1898-1918 (2007).

267	Ruprecht, E., Donath, T. W., Otte, A. & Eckstein, R. L. Chemical effects of a dominant grass on seed germination of four familial pairs of dry grassland species. *Seed Sci. Res.* **18**, 239-248 (2008).

268	Ruprecht, E., Jozsa, J., Olvedi, T. B. & Simon, J. Differential effects of several "litter" types on the germination of dry grassland species. *J. Veg. Sci.* **21**, 1069-1081 (2010).

269	Russo, V. M., Webber, C. L. & Myers, D. L. Kenaf extract affects germination and post-germination development of weed, grass and vegetable seeds. *Ind. Crop. Prod.* **6**, 59-69 (1997).

270	Ruwanza, S., Gaertner, M., Esler, K. J. & Richardson, D. M. Allelopathic effects of invasive Eucalyptus camaldulensis on germination and early growth of four native species in the Western Cape, South Africa. *South. Forests* **77**, 91-105 (2015).

271	Sajna, N. Habitat preference within its native range and allelopathy of garlic mustard Alliaria petiolata. *Pol. J. Ecol.* **65**, 46-56 (2017).

272	Samedani, B. *et al.* Phytotoxic effects of Pueraria javanica litter on growth of weeds Asystasia gangetica and Pennisetum polystachion. *Allelopathy J.* **32**, 191-202 (2013).

273	Sampietro, D. A., Sgariglia, M. A., Soberon, J. R., Quiroga, E. N. & Vattuone, M. A. Role of sugarcane straw allelochemicals in the growth suppression of arrowleaf sida. *Environ. Exp. Bot.* **60**, 495-503 (2007).

274	San Emeterio, L., Arroyo, A. & Canals, R. M. Allelopathic potential of Lolium rigidum Gaud. on the early growth of three associated pasture species. *Grass and Forage Science* **59**, 107-112 (2004).

275	Santos, P. C. *et al.* Phytotoxicity of Tagetes erecta L. and Tagetes patula L. on plant germination and growth. *S. Afr. J. Bot.* **100**, 114-121 (2015).

276	Sarika & Rao, P. B. Effect of weed extracts on germination, seedling growth and protein in french bean (Phaseolus vulgaris L.) varieties. *Allelopathy J.* **17**, 223-234 (2006).

277	Sarkar, E., Chatterjee, S. N. & Chakraborty, P. Allelopathic effect of Cassia tora on seed germination and growth of mustard. *Turk. J. Bot.* **36**, 488-494 (2012).

278	Saxena, M. K. Aqueous leachate of Lantana camara kills water hyacinth. *J. Chem. Ecol.* **26**, 2435-2447 (2000).

279	Sera, B. Effects of Soil Substrate Contaminated by Knotweed Leaves on Seed Development. *Pol. J. Environ. Stud.* **21**, 713-717 (2012).

280	Shafique, S., Bajwa, R. & Shafique, S. Tagetes Erectus L. - a Potential Resolution for Management of Parthenium Hysterophorus L. *Pak. J. Bot.* **43**, 885-894 (2011).

281	Shang, Z. H., Tang, Y. & Long, R. J. Allelopathic effect of Aconitum pendulum (Ranunculaceae) on seed germination and seedlings of five native grass species in the Tibetan Plateau. *Nord. J. Bot.* **29**, 488-494 (2011).

282	Shang, Z. H. & Xu, S. G. Allelopathic testing of Pedicularis kansuensis (Scrophulariaceae) on seed germination and seedling growth of two native grasses in the Tibetan plateau. *Phyton-Int. J. Exp. Bot.* **81**, 75-79 (2012).

283	Shanthi, R., Jahir Hussain, K. & Sanjayan, K. P. Effect of aqueous root and shoot extract of weeds on germination and seedling growth of cotton. *Allelopathy J.* **19**, 525-534 (2007).

284	Shao, H., Huang, X., Wei, X. & Zhang, C. Phytotoxic effects and a phytotoxin from the invasive plant Xanthium italicum Moretti. *Molecules* **17**, 4037-4046 (2012).

285	Shao, H., Huang, X., Zhang, Y. & Zhang, C. Main alkaloids of Peganum harmala L. and their different effects on dicot and monocot crops. *Molecules* **18**, 2623-2634 (2013).

286	Siddiqui, Z. S. Allelopathic effects of black pepper leachings on Vigna mungo (L.) Hepper. *Acta Physiol. Plant.* **29**, 303-308 (2007).

287	Siemens, T. J. & Blossey, B. An evaluation of mechanisms preventing growth and survival of two native species in invasive Bohemian knotweed (Fallopia xbohemica, Polygonaceae). *Am J Bot* **94**, 776-783 (2007).

288	Silva, E. R., Lazarotto, D. C., Schwambach, J., Overbeck, G. E. & Soares, G. L. G. Phytotoxic effects of extract and essential oil of Eucalyptus saligna (Myrtaceae) leaf litter on grassland species. *Aust. J. Bot.* **65**, 172-182 (2017).

289	Silva, E. R., Overbeck, G. E. & Soares, G. L. G. Phytotoxicity of volatiles from fresh and dry leaves of two Asteraceae shrubs: Evaluation of seasonal effects. *S. Afr. J. Bot.* **93**, 14-18 (2014).

290	Silva, R. M. G., Brante, R. T., Santos, V. H. M., Mecina, G. F. & Silva, L. P. Phytotoxicity of ethanolic extract of turnip leaves (Raphanus sativus l.). *Biosci. J.* **30**, 891-902 (2014).

291	Silva, R. M. G., Livio, A. A., Santos, V. H. M., Mecina, G. F. & Silva, L. P. Allelopathy and phytotoxicity of Zanthoxylum rhoifolium Lam. *Allelopathy J.* **30**, 221-234 (2012).

292	Singh, A., Singh, D. & Singh, N. B. Allelochemical stress produced by aqueous leachate of Nicotiana plumbaginifolia Viv. *Plant Growth Regul.* **58**, 163-171 (2009).

293	Singh, B., Uniyal, A. K. & Todaria, N. P. Studies on Allelopathic Influence ofZanthoxylum armatumD.C. on Important Field Crops Seeking Its Sustainable Domestication in Existing Agroforestry Systems of Garhwal Himalaya, India. *J. Sustain. Agric.* **30**, 87-95 (2007).

294	Singh, H. P. *et al.* Negative effect of litter of invasive weed Lantana camara on structure and composition of vegetation in the lower Siwalik Hills, northern India. *Environ Monit Assess* **186**, 3379-3389 (2014).

295	Singh, N. B., Pandey, B. N. & Singh, A. Allelopathic effects of Cyperus rotundus extract in vitro and ex vitro on banana. *Acta Physiol. Plant.* **31**, 633-638 (2009).

296	Singh, N. B., Singh, A. & Singh, D. Autotoxic effects of Lycopersicon esculentum. *Allelopathy J.* **22**, 429-442 (2008).

297	Skrzypek, E., Repka, P., Stachurska-Swakon, A., Barabasz-Krasny, B. & Mozdzen, K. Allelopathic Effect of Aqueous Extracts from the Leaves of Peppermint (Mentha x piperita L.) on Selected Physiological Processes of Common Sunflower (Helianthus annuus L.). *Not. Bot. Horti Agrobot. Cluj-Na.* **43**, 335-342 (2015).

298	Small, C. J., White, D. C. & Hargbol, B. Allelopathic influences of the invasive Ailanthus altissima on a native and a non-native herb1,2. *The Journal of the Torrey Botanical Society* **137**, 366-372 (2010).

299	Snyman, H. A. Allelopathic potential, seed ecology and germination of the encroacher shrubSeriphium plumosum. *African Journal of Range & Forage Science* **27**, 29-37 (2010).

300	Sousa, F. G., Denardin, R. B. N., Moura, N. F. & Drevs, S. Allelopathy and genotoxic potential of Casearia sylvestris Sw. Extracts. *Allelopathy J.* **20**, 195-202 (2007).

301	Souza, F. M., Gandolfi, S., de Andrade Perez, S. C. J. G. & Rodrigues, R. R. Allelopathic potential of bark and leaves of Esenbeckia leiocarpa Engl. (Rutaceae). *Acta Bot. Bras.* **24**, 169-174 (2010).

302	Souza-Alonso, P., Gonzalez, L. & Cavaleiro, C. Ambient has become strained. Identification of Acacia dealbata Link volatiles interfering with germination and early growth of native species. *J Chem Ecol* **40**, 1051-1061 (2014).

303	Suksungworn, R. *et al.* Phytotoxic effect of Haldina cordifolia on germination, seedling growth and root cell viability of weeds and crop plants. *NJAS-Wagen. J. Life Sci.* **78**, 175-181 (2016).

304	Suman, A., Shahi, H. N., Singh, P. & Gaur, A. Allelopathic influence of Vigna mungo (black gram) seeds on germination and radical growth of some crop plants. *Plant Growth Regul.* **38**, 69–74 (2002).

305	Sun, H. & Wang, Y. Potential allelopathic effects of allelochemicals in aqueous extracts of leaves and root exudates of Capsicum annuum on vegetable crops. *Allelopathy J.* **35**, 11-22 (2015).

306	Sun, J. F. *et al.* Allelopathic effects of Solidago canadensis L. *Allelopathy J.* **41**, 189-200 (2017).

307	Sunar, S., Aksakal, O., Yildirim, N. & Agar, G. Determination of the Genotoxic Effects of Verbascum speciosum Schrad. Extracts on Corn (Zea mays L.) Seeds. *Rom. Biotech. Lett.* **14**, 4820-4826 (2009).

308	Sunmonu, T. O. & Van Staden, J. Phytotoxicity evaluation of six fast-growing tree species in South Africa. *S. Afr. J. Bot.* **90**, 101-106 (2014).

309	Tang, S. C., Pan, Y. M., Wei, C. Q., Liu, M. C. & Cen, Y. X. Allelopathic interactions between the invasive weed Chromolaena odorata and native plant Vitex negundo. *Allelopathy J.* **30**, 209-220 (2012).

310	Tanveer, A. *et al.* Assessing the potential of the water soluble allelopaths of Marsilea minuta in rice and wheat. *Planta Daninha* **33**, 231-239 (2015).

311	Terzi, I., Kocacaliskan, I. & Demir, Y. Allelopathic Effects of Some Tree Leaf Extracts on Seed Germination and Seedling Growth of Turf Grasses. *J. Environ. Prot. Ecol.* **14**, 1236-1243 (2013).

312	Tesio, F., Vidotto, F. & Ferrero, A. Allelopathic persistence of Helianthus tuberosus L. residues in the soil. *Sci. Hortic.* **135**, 98-105 (2012).

313	Tian, Y. H., Feng, Y. L. & Liu, C. Addition of activated charcoal to soil after clearing Ageratina adenophora stimulates growth of forbs and grasses in China. *Tropical Grasslands* **41**, 285-291 (2007).

314	Tinnin, R. O. & Muller, C. H. The Allelopathic Influence of Avena fatua: The Allelopathic Mechanism. *Bulletin of the Torrey Botanical Club* **99**, 287 (1972).

315	Tomar, N. S., Sharma, M. & Agarwal, R. M. Phytochemical analysis of Jatropha curcas L. during different seasons and developmental stages and seedling growth of wheat (Triticum aestivum L) as affected by extracts/leachates of Jatropha curcas L. *Physiol Mol Biol Plants* **21**, 83-92 (2015).

316	Travlos, I. S. & Paspatis, E. A. Allelopathic effects of Heliotrope (Helitropium europaeum L.) on Avena sativa, Phaseolus vulgaris and Spirodela polvrhiza. *Allelopathy J.* **21**, 397-404 (2008).

317	Tseng, M.-H., Kuo, Y.-H., Chen, Y.-M. & Chou, C.-H. Allelopathic potential of Macaranga tanarius (l.) muell.–arg. *J. Chem. Ecol.* **29**, 1269-2186 (2003).

318	Uddin, M. N., Caridi, D. & Robinson, R. W. Phytotoxic evaluation of Phragmites australis: an investigation of aqueous extracts of different organs. *Mar. Freshw. Res.* **63**, 777-787 (2012).

319	Uddin, M. N., Robinson, R. W., Buultjens, A., Al Harun, M. A. Y. & Shampa, S. H. Role of allelopathy of Phragmites australis in its invasion processes. *J. Exp. Mar. Biol. Ecol.* **486**, 237-244 (2017).

320	Uddin, M. N., Robinson, R. W., Caridi, D. & Al Harun, M. A. Suppression of native Melaleuca ericifolia by the invasive Phragmites australis through allelopathic root exudates. *Am J Bot* **101**, 479-487 (2014).

321	Uddin, M. N., Robinson, R. W., Caridi, D. & Harun, M. A. Y. Is phytotoxicity of Phragmites australis residue influenced by decomposition condition, time and density? *Mar. Freshw. Res.* **65**, 505-516 (2014).

322	van de Voorde, T. F. J., Ruijten, M., van der Putten, W. H. & Bezemer, T. M. Can the negative plant–soil feedback of Jacobaea vulgaris be explained by autotoxicity? *Basic Appl. Ecol.* **13**, 533-541 (2012).

323	Vance, H. D. & Francko, D. A. Allelopathic potential of Nelumbo lutea(willd.) pers. To alter growth of Myriophyllum spicatuml and Potamogeton pectinatusl. *J. Freshw. Ecol.* **12**, 405-409 (1997).

324	Vázquez-de-Aldana, B. R., Romo, M., García-Ciudad, A., Petisco, C. & García-Criado, B. Infection with the fungal endophyte Epichloë festucae may alter the allelopathic potential of red fescue. *Ann. Appl. Biol.* **159**, 281-290 (2011).

325	Velicka, R. *et al.* Allelopathic effects of aqueous extracts of rape residues on winter wheat seed germination and early growth. *J. Food Agric. Environ.* **10**, 1053-1057 (2012).

326	Viard-Cretat, F., Baptist, F., Secher-Fromell, H. & Gallet, C. The allelopathic effects of Festuca paniculata depend on competition in subalpine grasslands. *Plant Ecol.* **213**, 1963-1973 (2012).

327	Viard-Cretat, F., Gallet, C., Lefebvre, M. & Lavorel, S. A leachate a day keeps the seedlings away: mowing and the inhibitory effects of Festuca paniculata in subalpine grasslands. *Ann Bot* **103**, 1271-1278 (2009).

328	Vidotto, F., Tesio, F. & Ferrero, A. Allelopathic effects of Ambrosia artemisiifolia L. in the invasive process. *Crop Prot.* **54**, 161-167 (2013).

329	Viles, A. L. & Reese, R. N. Allelopathic potential of Echinacea Angustifolia D.C. *Environ. Exp. Bot.* **35**, 39-43 (1996).

330	Wang, C. *et al.* The allelopathic effects of invasive plant Solidago canadensis on seed germination and growth of Lactuca sativa enhanced by different types of acid deposition. *Ecotoxicology* **25**, 555-562 (2016).

331	Wang, C. Y., Liu, J., Xiao, H. G., Zhou, J. W. & Du, D. L. Nitrogen deposition influences the allelopathic effect of an invasive plant on the reproduction of a native plant: Solidago canadensis versus Pterocypsela laciniata. *Pol. J. Ecol.* **65**, 87-96 (2017).

332	Wang, J. C. *et al.* Allelopathic effects of Jatropha curcas on marigold (Tagetes erecta L.). *Allelopathy J.* **24**, 123-130 (2009).

333	Wang, L., Deng, D. Y., Tang, L., Yuan, K. & Li, Y. X. in *Proceedings of the 2015 International Conference on Industrial Technology and Management Science* Vol. 34 *ACSR-Advances in Comptuer Science Research* (ed J. Zhao) 1023-1026 (Atlantis Press, 2015).

334	Wang, P., Liang, W., Kong, C. & Jiang, y. Allelopathic potentials of volatile allelochemicals of Ambrosia trifida L. on other plants. *Allelopathy J.* **15**, 131-136 (2005).

335	Wang, R. *et al.* Allelopathic potential of Tephrosia vogelii Hook. f.: Laboratory and field evaluation. *Allelopathy J.* **28**, 53-62 (2011).

336	Wang, X. F., Hassani, D., Cheng, Z. W., Wang, C. Y. & Wu, J. Allelopathy of the invasive plant Bidens frondosa on the seed germination of Geum japonicum var. chinense. *Genet Mol Res* **13**, 10592-10598 (2014).

337	Wang, Y. *et al.* Allelopathic effects of corn and soybean root leachates on tuber sprouting and plantlet growth of potato. *Allelopathy J.* **31**, 117-128 (2013).

338	Watanabe, Y. *et al.* Assessment of allelopathic activities in female and male individuals of asparagus seedlings and regenerants. *J. Japan. Soc. Hort. Sci.* **80**, 169–174 (2011).

339	Weißhuhn, K. & Prati, D. Activated carbon may have undesired side effects for testing allelopathy in invasive plants. *Basic Appl. Ecol.* **10**, 500-507 (2009).

340	Wille, W., Thiele, J., Walker, E. A. & Kollmann, J. Limited evidence for allelopathic effects of giant hogweed on germination of native herbs. *Seed Sci. Res.* **23**, 157-162 (2013).

341	Wixted, K. L. & McGraw, J. B. Competitive and allelopathic effects of garlic mustard (Alliaria petiolata) on American ginseng (Panax quinquefolius). *Plant Ecol.* **208**, 347-357 (2009).

342	Wu, A. P., Huang, Z., Miao, S. L. & Dong, M. Effects of Mikania micrantha extracts and their exposure time on seed vigour, seed germination and seedling growth of plants. *Allelopathy J.* **25**, 503-512 (2010).

343	Wu, A.-P. *et al.* Differential belowground allelopathic effects of leaf and root of Mikania micrantha. *Trees* **23**, 11-17 (2008).

344	Wu, G. L., Ren, G. H. & Shi, Z. H. Phytotoxic effects of a dominant weed Ligularia virgaurea on seed germination of Bromus inermis in an alpine meadow community. *Plant Ecol. Evol.* **144**, 275-280 (2011).

345	Wu, J. R., Peng, S. L., Zhao, H. B. & Xiao, H. L. Allelopathic effects of *Wedelia trilobata* residues on lettuce germination and seedling growth. *Allelopathy J.* **22**, 197-203 (2008).

346	Wu, L., Guo, X. & Harivandi, M. A. Allelopathic effects of phenolic acids detected in buffalograss (Buchloe dactyloides) clippings on growth of annual bluegrass (Poa annua) and buffalograss seedlings. *Environ. Exp. Bot.* **39** 159–167 (1998).

347	Wurst, S., Vender, V. & Rillig, M. C. Testing for allelopathic effects in plant competition: does activated carbon disrupt plant symbioses? *Plant Ecol.* **211**, 19-26 (2010).

348	Wyse, S. V. & Burns, B. R. Effects of Agathis australis (New Zealand kauri) leaf litter on germination and seedling growth differs among plant species. *N. Z. J. Ecol.* **37**, 178-183 (2013).

349	Xiao, C. L. *et al.* Autotoxic effects of root exudates of soybean. *Allelopathy J.* **18**, 121-128 (2006).

350	Xie, W. Y., Hu, N. Y., Yu, Y. N., Niu, Z. J. & Li, X. Resistance of plant community to exotic invasive Mikania micrantha through allelopathy of multiple species. *Allelopathy J.* **41**, 239-248 (2017).

351	Xu, N. *et al.* Composition of Welsh onion (Allium fistulosum L.) root exudates and their allelopathy on cucumber sprouts and Fusarium oxysporum f.sp. cucumerinum. *Allelopathy J.* **32**, 243-256 (2013).

352	Xuan, T. D., Tsuzuki, E., Terao, H., Matsuo, M. & Khanh, T. D. Correlation between growth inhibitory exhibition and suspected allelochemicals (phenolic compounds) in the extract of alfalfa (Medicago sativa L.). *Plant Production Science* **6**, 165-171 (2003).

353	Yang, G. Q., Qiu, W. R., Jin, Y. N. & Wan, F. H. Potential allelochemicals from root exudates of invasive Ageratina adenophora. *Allelopathy J.* **32**, 233-242 (2013).

354	Yang, G. Q., Wan, F. H., Liu, W. X. & Zhang, X. W. Physioligical effects of allelochemicals from leachates of Ageratina adenophora (Spreng.) on rice seedlings. *Allelopathy J.* **18**, 237-246 (2006).

355	Yang, L., Chen, Y., Huang, Y., Wang, J. & Wen, M. Mixed allelopathic effect of eucalyptus leaf litter and understorey fern in south china. *J. Trop. For. Sci.* **28**, 436-445 (2016).

356	Yang, R. X., Mei, L. X., Tang, J. J. & Chen, X. Allelopathic effects of invasive Solidago canadensis L. on germination and growth of native Chinese plant species. *Allelopathy J.* **19**, 241-248 (2007).

357	Yang, X. *et al.* The extraction, isolation and identification of exudates from the roots of Flaveria bidentis. *J. Integr. Agric.* **13**, 105-114 (2014).

358	Young, G. P. & Bush, J. K. Assessment of the allelopathic potential of Juniperus ashei on germination and growth of Bouteloua curtipendula. *J Chem Ecol* **35**, 74-80 (2009).

359	Yu, F., Ma, Y., Wei, G. & Zhao, S. Allelopathic potential of Astragalus adsurgens Pall on the growth of cultured Stelleria chamaejasme L. *Allelopathy J.* **17**, 255-264 (2006).

360	Yuan, Y. G. *et al.* Enhanced allelopathy and competitive ability of invasive plant Solidago canadensis in its introduced range. *J. Plant Ecol.* **6**, 253-263 (2013).

361	Yun, K. W. & Maun, M. A. Allelopathic potential of Artemisia campestris ssp. caudata on Lake Huron sand dunes. *Canadian Journal of Botany* **75**, 1903- 1912 (1997).

362	Zasada, I. A., Klassen, W., Meyer, S. L., Codallo, M. & Abdul-Baki, A. A. Velvetbean (Mucuna pruriens) extracts: impact on Meloidogyne incognita survival and on Lycopersicon esculentum and Lactuca sativa germination and growth. *Pest Manag Sci* **62**, 1122-1127 (2006).

363	Zhang, C. L. & Fu, S. L. Allelopathic effects of eucalyptus and the establishment of mixed stands of eucalyptus and native species. *For. Ecol. Manage.* **258**, 1391-1396 (2009).

364	Zhang, F. J., Guo, J. Y., Chen, F. X., Guo, A. Y. & Wan, F. H. Assessment of allelopathic effects of residues of Flaveria bidentis (L.) Kuntze on wheat seedlings. *Arch. Agron. Soil Sci.* **58**, 257-265 (2012).

365	Zhang, F. J., Guo, J. Y., Chen, F. X., Liu, W. X. & Wan, F. H. Identification of Volatile Compounds Released by Leaves of the Invasive Plant Croftonweed (Ageratina adenophora, Compositae), and their Inhibition of Rice Seedling Growth. *Weed Sci.* **60**, 205-211 (2012).

366	Zhang, H. Y. *et al.* Allelopathic potential of flavonoids identified from invasive plant Conyza canadensis on Agrostis stolonifera and Lactuca sativa. *Allelopathy J.* **41**, 223-238 (2017).

367	Zhang, M., Ling, B., Kondo, C., Liang, G. & Dong, Y. Allelopathic effects of lantana (Lantana camera L.) on water hyacinth (Eichhornia crassipes (Mart.) Solms). *Allelopathy J.* **15**, 125-130 (2005).

368	Zhang, Q., Peng, S. & Zhang, Y. Allelopathic potential of reproductive organs of exotic weed Lantana camara. *Allelopathy J.* **23**, 213-220 (2009).

369	Zhang, S., Liu, J., Bao, X. & Niu, K. Seed-to-seed potential allelopathic effects between Ligularia virgaurea and native grass species of Tibetan alpine grasslands. *Ecol. Res.* **26**, 47-52 (2010).

370	Zhang, Y. J. *et al.* The allelopathic effect of Potentilla acaulis on the changes of plant community in grassland, northern China. *Ecol. Res.* **30**, 41-47 (2015).

371	Zhang, Z. H., Hu, B. Q. & Hu, G. in *Environmental Protection and Resources Exploitation, Pts 1-3* Vol. 807-809 *Advanced Materials Research* (eds Z. Liu, X. Dong, Z. Liu, & Q. Liu) 719-+ (Trans Tech Publications Ltd, 2013).

372	Zhao, H. & Peng, S. Activity of Mikania micrantha H.B.K. allelochemicals released in soil. *Allelopathy J.* **24**, 405-412 (2009).

373	Zhao, Y. J., Wang, Y., Shao, D., Yang, J. & Liu, D. Autotoxicity of panax quinquefolium l. *Allelopathy J.* **15**, 67-74 (2005).

374	Zhu, B. & Georgian, S. E. Interactions between invasive Eurasian watermilfoil and native water stargrass in Cayuga Lake, NY, USA. *J. Plant Ecol.* **7**, 499-508 (2014).

375	Zhu, H. & Mallik, A. U. Interactions between *Kalmia* and black spruce: Isolation and identification of allelopathic compounds. *J. Chem. Ecol.* **20**, 407-421 (1994).

376	Zhu, W., Liu, J. W., Ye, J. L. & Li, G. H. Effects of phytotoxic extracts from peach root bark and benzoic acid on peach seedlings growth, photosynthesis, antioxidance and ultrastructure properties. *Sci. Hortic.* **215**, 49-58 (2017).

377	Zhu, X., Zhang, J. & Ma, K. Soil biota reduce allelopathic effects of the invasive Eupatorium adenophorum. *PLoS One* **6**, e25393 (2011).

378	Zuo, S., Ma, Y., Deng, X. & Li, X. Allelopathy in wheat genotypes during the germination and seedling stages. *Allelopathy J.* **15**, 21-30 (2005).
